# Inactivation mechanism and efficacy of grape seed extract for Human Norovirus surrogate

**DOI:** 10.1101/2021.12.03.471102

**Authors:** Chamteut Oh, Ratul Chowdhury, Laxmicharan Samineni, Joanna L Shisler, Manish Kumar, Thanh H. Nguyen

## Abstract

Proper disinfection of harvested food and water is critical to minimize infectious disease. Grape seed extract (GSE), a commonly used health supplement, is a mixture of plant-derived polyphenols. Polyphenols possess anti-microbial and -fungal properties, but anti-viral effects are not well-known. Here we show that GSE outperformed chemical disinfectants (e.g., free chlorine and peracetic acids) in inactivating Tulane virus, a human norovirus surrogate. GSE induced virus aggregation, an event that correlated with a decrease in virus titers. This aggregation and disinfection was not reversible. Molecular docking simulations indicate that polyphenols potentially formed hydrogen bonds and strong hydrophobic interactions with specific residues in viral capsid proteins. Together, these data suggest that polyphenols physically associate with viral capsid proteins to aggregate viruses as a means to inhibit virus entry into the host cell. Plant-based polyphenols like GSE are an attractive alternative to chemical disinfectants to remove infectious viruses from water or food.

**Importance:** Human noroviruses are major food- and water-borne pathogens, causing approximately 20% of all cases of acute gastroenteritis cases in developing and developed countries. Proper sanitation or disinfection are critical strategies to minimize human norovirus-caused disease until a reliable vaccine is created. Grape seed extract (GSE) is a mixture of plant-derived polyphenols that is used as a health supplement. Polyphenols are known for antimicrobial, antifungal, and antibiofilm activities, but antiviral effects are not well-known. In studies here, plant-derived polyphenols outperformed chemical disinfectants (e.g., free chlorine and peracetic acids) in inactivating Tulane virus, a human norovirus surrogate. Based on data from additional molecular assays and molecular docking simulations, the current model is that the polyphenols in GSE bind to the Tulane virus capsid, an event that triggers virion aggregation. It is thought that this aggregation prevents Tulane virus from entering host cells.

## Introduction

Human noroviruses cause approximately 20% of all cases of acute gastroenteritis in developing and developed countries (1). In the United States, human noroviruses cause about 5.5 million cases of food borne illnesses per year, and about 2 billion dollars in economic loss (2, 3). Noroviruses are transmitted primarily by the fecal-oral route, including ingestion of contaminated food and water or via person-to-person contacts (4). Thus, inactivating human noroviruses present in contaminated food or water is important.

Sodium hypochlorite (NaClO) is regarded as the cheapest and most effective disinfectant to inactivate noroviruses on produce surfaces (5), stainless steel surfaces (6), or in liquid solutions (7). Fresh and fresh-cut produce are generally sanitized with residual chlorine concentrations of 50-200 μg/mL (8). Unfortunately, NaClO is toxic, and produces carcinogenic disinfection by-products (9–11). Thus, both consumers and industry are seeking methods to naturally or minimally process foods with minimal/zero chemical additives (12, 13).

Plants and fruits synthesize polyphenols, chemicals that protect them against damage from external stresses such as infection. Grape seed extract (GSE) is a by-product of grape juice and wine production, and is mass-produced at an affordable price (14, 15). GSE already is sold as an FDA-approved health supplement (16). GSE polyphenols have antimicrobial effects (17) and are considered popular alternatives to chemical disinfectants. GSE possesses anti-viral activity against feline calicivirus (FCV-F9), murine norovirus (MNV-1), bacteriophage MS2, and hepatitis A virus (14, 18, 19). However, the mechanism of virus inactivation is unclear (16). This information is critical because, if GSE is to be used by industries as a natural disinfectant, then researchers must understand how GSE inactivates different types of viruses.

The objective of this study was to examine GSE-induced inactivation of Tulane virus, a surrogate for human norovirus. Tulane virus is an ideal surrogate for norovirus because of its structural similarities to human noroviruses (20–22). Thus, Tulane virus has been used to provide more information about potential inactivation of norovirus (7, 23–27). Once GSE disinfection was established, we identified the mechanism of this inactivation using both molecular assays and computer modeling. Studies showed that GSE’s main disinfection action occurred due to virus aggregation. Molecular docking simulations identified potential interactions between GSE polyphenols and viral capsids. This study is the first observation that shows plant-derived polyphenols can outperform chemical disinfectants in safely and sustainably controlling water-borne and food-borne pathogens, and also provides the first mechanism for GSE-induced virus inactivation. Thus, GSE may be an attractive disinfectant for industry because of its safety, efficacy and lower impact on the environment.

## Results

### GSE outperforms chemical disinfectants by 3-log_10_ TV titer reduction

**Fig. 1** shows results from the examination of disinfection properties of GSE against Tulane virus, using a range of different GSE concentrations, Tulane virus (TV) concentrations, and incubation times. For all experiments, GSE was incubated with purified TV. GSE activity was halted by the addition of FBS, and plaque-forming units (PFUs) were quantified as a measure of virus inactivation. We tested GSE concentrations ranging from 42 μg/mL to 678 μg/mL because similar GSE concentrations were shown to inactivate other enteroviruses (14, 18, 19).

**Fig. 1.**
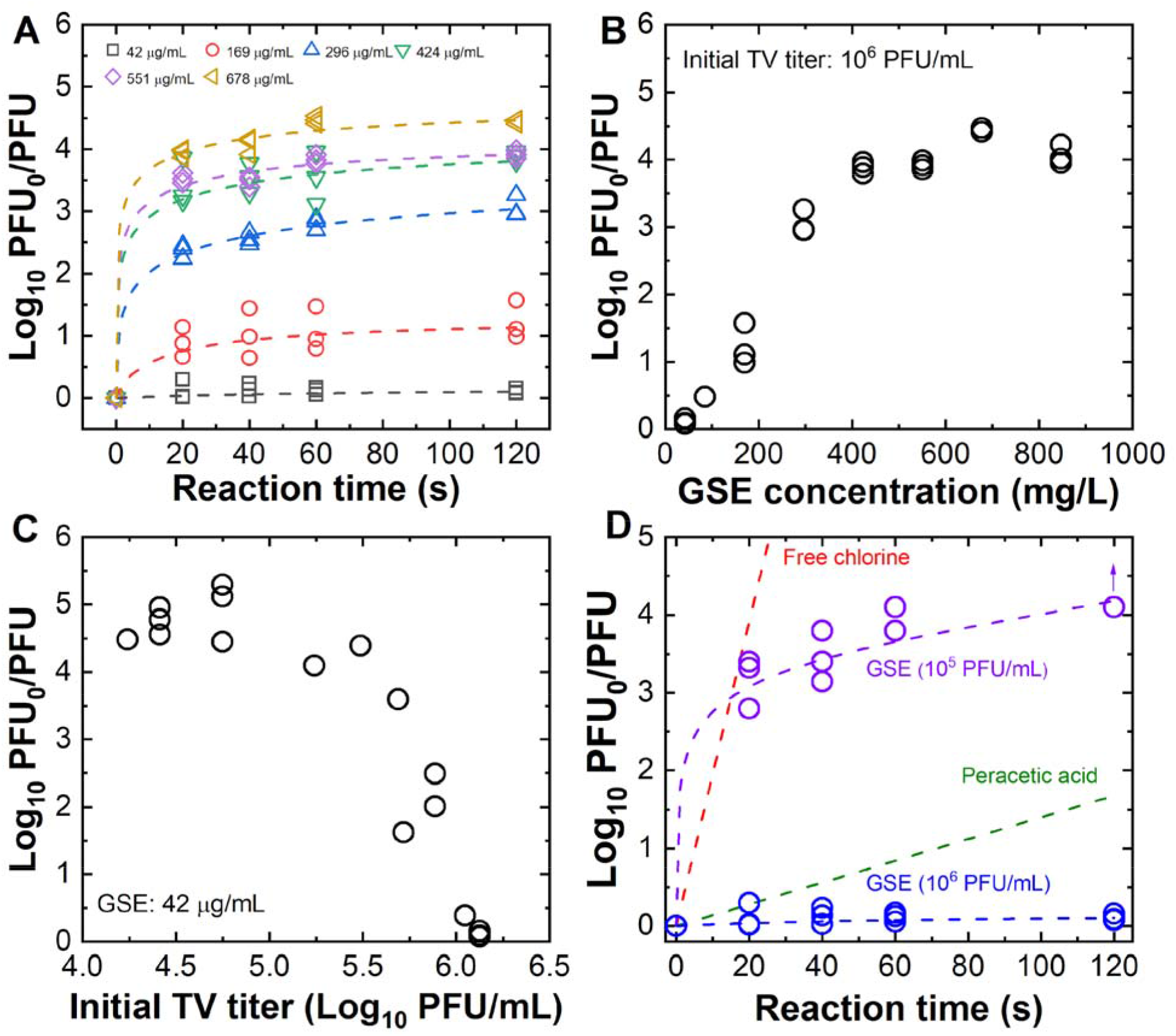
Inactivation effects of GSE on Tulane virus. 10^6^ PFU Tulane virus was incubated with the indicated amount of GSE. FBS was added to quench each reaction at the indicated times. Initial virus titer (PFU_0_) was divided by the virus titer at each indicated time (PFU). Thus, the y-axis (Log_10_ PFU_0_/PFU) indicates the decrease in virus titer caused by GSE on log_10_ scale. For all experiments, each symbol represents one molecular replicate. **(A)** The dashed lines were trend lines calculated by the pseudo-second-order model. TV inactivation as a function of GSE concentration and time. **(B)** TV inactivation as a function of GSE concentration. In this experiment 10^6^ PFU TV was incubated for 120 seconds with the indicated GSE concentrations. **(C)** TV inactivation as a function of GSE concentration. Varying amounts of TV (indicated on the x-axis) were added to reactions containing 42 μg/mL GSE. All reactions were quenched with FBS after a 120 second incubation. **(D)** A comparison of GSE versus peracetic acid and free chlorine inactivation. 10^5^ or 10^6^ PFU TV was incubated with GSE (42 μg/mL) for 120 sec. The trend lines for GSE were determined by the pseudo-second-order model. The arrow indicates a detection limit. Trend lines for 42 μg /mL free chlorine or peracetic acid were derived by Chick’s law with the rate constants determined by our previous studies(7, 27).

TV inactivation increased when GSE concentration and duration of incubation increased (**Fig. 1A**). The inactivation curves were fitted to both the pseudo-second-order model and Chick’s law. The correlation coefficients (R^2^ values), which reflect the goodness of fit, obtained by the pseudo-second-order model (0.99 to 1.00) were higher than those by the Chick’s law (0.34 to 0.54) except for the lowest GSE concentration (42 μg/mL) where no significant reduction in virus titers was detected (one-way ANOVA, p>0.05). Parameters from the inactivation kinetics experiments are also listed in **Supplementary Table 1**. The fitting to the pseudo-second-order model showed that it took less than 120 seconds to reach a 95% of log_10_ PFU reductions at equilibrium state (i.e., t_95_<120 seconds) for all tested conditions except for the 42 μg/mL case.

In **Fig. 1A**, better fitting achieved by the pseudo-second-order model suggested that chemisorption is the dominant reaction between the polyphenols and virus particles (28–30). This finding suggested there were optimal GSE:virion ratios for disinfection, as supported by data in **Fig. 1B** and **1C. Fig. 1B** showed that TV inactivation increased as GSE concentrations increased, with GSE efficacy reaching a plateau at 424 μg/mL. This plateau in GSE efficacy shows a maximal number of GSE binding sites on TVs because GSE disinfection could be outcompeted by the addition of TV. **Fig. 1C** showed that GSE disinfection efficiency decreased when more TVs were added to a reaction, supporting the hypothesis that chemisorption occurs between GSE and TV. Interestingly, while 42 mg/mL GSE showed no disinfection activity for 10^6^ PFU TV (in **Fig. 1A**), it does indeed possess anti-viral properties when lower numbers of TV are present, further supporting the model that GSE polyphenols associate with TV.

We compared GSE inactivation rates to those of two commercially available chemical disinfectants (free chlorine and peracetic acid) based on Chick’s law and the rate constants obtained from a previous study (27) (**Fig. 1D**). GSE inactivated TV to a 3-log_10_ virus titer reduction within 16 seconds, a time identical to that of inactivation by free chlorine. In contrast, peracetic acid took much longer (211 seconds) to achieve the same levels of inactivation.

### GSE causes viral aggregation, and this is likely the main mechanism of GSE-induced virus inactivation

**Fig. 1** data showed that GSE inactivated TV, and data suggested that there was chemisorption of GSE to virions. Thus, one possibility is that GSE binds directly to TV to prevent virus-host interactions. **Fig. 2A** quantifies GSE polyphenols in the absence or presence of TV as a means to examine if GSE indeed binds to TV. We incubated 847 μg/mL GSE with 10^6^ PFU TV. The 2 mL mixture then subjected to ultracentrifugation to separate free versus virus-bound GSE, and GSE in supernatants of samples were quantified. As shown in **Fig. 2A**, there was a statistically significant lower GSE concentration in supernatants when TV was present, implying that GSE indeed binds to TV. Next, we used a light scattering analyzer (DelsaMax Pro, Beckman Coulter) to examine particle size distributions of intact TV and GSE (**Fig. 2B)**. Untreated TV showed a polydispersed multimodal size distribution with a major peak at 97 nm and two relatively smaller peaks at 309 and 1755 nm. A single TV virion has a diameter of 40 nm (31), thus most of the TVs in solution were present as dimers, and some populations of viruses were present as timers. Virus populations at 1755 nm were likely multi-virus aggregates. When TV was incubated with 15 μg /ml GSE, there was a shift in the size of the TV peak at 94 nm to approximately 300 nm and 1000 nm, implying that this concentration of GSE causes aggregation of TV into trimers and perhaps 10-mers. Experiments using higher concentrations of GSE resulted in greater shifts in sizes, suggesting that GSE is causing TV aggregation. GSE showed a strong single peak at around 1000 nm regardless of GSE concentrations. This peak is believed to represent insoluble polyphenols that are self-aggregated (32). When the GSE solutions were filtered with 0.1 μm filter, the monodispersed peak disappeared from the particle size distribution.

**Fig. 2.**
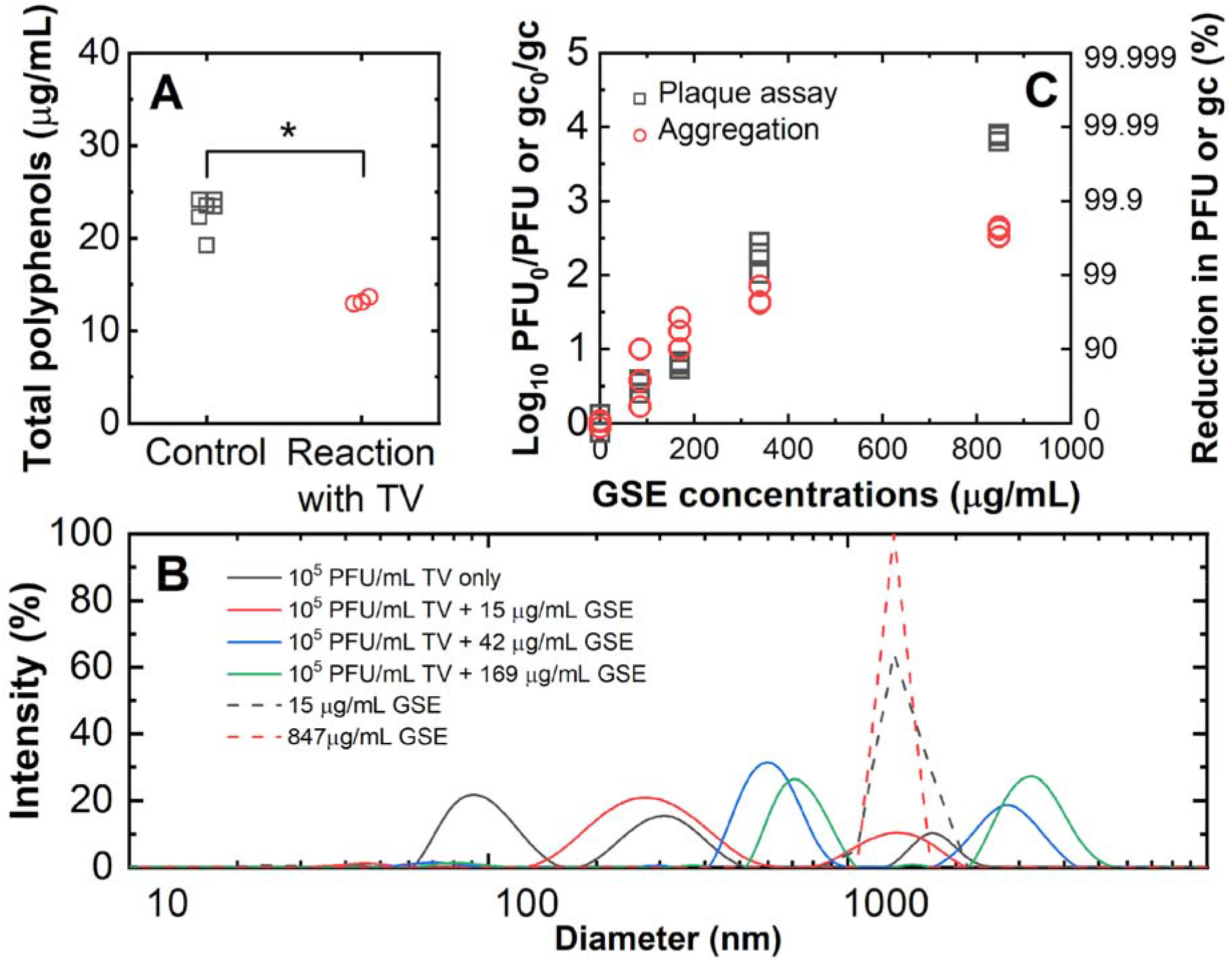
GSE-induced TV aggregation. **(A)** Solutions either lacking (control) or containing 10^6^ PFU TV were each incubated with 847 μg/mL of GSE followed by ultracentrifugation (150700 g) for 3 hours. GSE concentrations were quantified and statistical analysis was performed using the Mann-Whitney test (P<0.05) **(B)** The particle size distribution of TV in the presence or absence of GSE. **(C)** A dose-response curve for plaque and aggregation assays. Plaque assay results were normalized to the initial virus titer and presented in Log_10_ PFU_0_/PFU to indicate the decrease in virus titer with different total polyphenol concentrations. Aggregation assay was analyzed by the one-step RT-qPCR and presented in the normalized gene copy number (Log_10_ gc_0_/gc) to show the reduction in the number of virions. Equivalent reductions in PFU or gc (%) were also presented on the right y-axis for plaque assay and aggregation assay, respectively. For these experiments, 10^6^ PFU TV was incubated with varying concentrations of GSE for 2 minutes followed by quenching the reaction with FBS. Each symbol indicates one separate experiment where one plaque assay or three qPCR analyses were conducted.

To further test if GSE induces virus aggregation, virus aggregation and virus titers were quantified in parallel to compare the numbers of virus aggregates and inactivated virus particles after the GSE treatment. For assays shown in **Fig. 2C**, 10^6^ PFU TV was incubated with the indicated concentrations of GSE (85 to 847 μg/mL) for 2 minutes followed by quenching the reaction with FBS. Viruses were then either quantified by plaque assay or tested using an aggregation assay. Aggregation assays were designed as follows: untreated or GSE-treated TV were subjected to filtration using a 0.1 μm sized pore filter. Because a single TV virion has a diameter of 40 nm (31), we expect that single and dimeric TV would easily filter through the pore while virus aggregates with three or more virions would be trapped in the filter. Note that our aggregation assay cannot determine a specific number of virions in virus aggregates that can pass through the filter. Viruses that passed through the filter were quantified by one-step RT-qPCR. The ratio of genome copies for untreated versus treated viruses (Log_10_ gc_0_/gc) were plotted on the y-axis in **Fig. 2C**. Similar to **Fig. 1**, plaque assays measured virus titers in untreated samples (PFU_0_) versus GSE-treated samples (PFU), and data are presented as the ratio of virus titers before and after exposure to GSE (Log_10_ PFU_0_/PFU) on the y-axis in **Fig. 2C**. Virus aggregates and plaque were very similar when GSE concentrations below 400 ng/ml were used. However, virus aggregates were significantly lower than inactivated virus titers when 847 μg/mL GSE was present. This discrepancy shows the number of virus particles that are larger than 0.1 μm does not fully explain the number of inactivated virus particles determined by the plaque assay. Nevertheless, paired t-test revealed that there was no significant difference between the plaque assay and aggregation assay results over the different GSE concentrations (P>0.05). Thus, our current model is that GSE interacts with TV capsid, causing TV aggregation. In turn, aggregation would likely prevent TV from entering host cells. We also demonstrate that GSE aggregation was not reversible; removal of GSE from TV-containing solutions did not make aggregated TV return back into single particle solutions (**Supplementary Text 1**).

### Molecular docking indicated polyphenols strongly bound to capsid proteins

Data from **Fig. 2** indicated that GSE is physically associated with TV and induced virus aggregation, and that this event likely prevents TV from entering the host cell to begin the virus life cycle. To further investigate this possibility, we conducted molecular docking simulations, an approach that was used to identify interactions between polyphenolic compounds and proteins (33, 34). GSE is a mixture of various polyphenolic compounds (35, 36). We selected nine polyphenolic compounds present in GSE for molecular docking experiments. We chose these compounds based on abundance (36), and this included two phenolic acids (gallic acid and protocatechuic acid), three monomer flavan-3-ols (catechin, epicatechin, and epicatechin gallate), and four dimer flavan-3-ols based on LC-MS analysis (procyanidin B1, B2, B3, and B4) (**Supplementary Table 2**). Although we conducted inactivation experiments using TV, we could not perform molecular docking simulations with the TV capsid because there is no entry for the TV capsid in the Protein Data Bank (PDB, https://www.rcsb.org/). Cryo-EM analysis has shown similarity between the TV and human norovirus (HuNoV) capsids (31), and the HuNoV capsid structure is present in the PBD. Thus, we studied the interaction of the GSE polyphenols with the HuNoV VP1 capsid proteins to understand how GSE may interact with similar TV capsid proteins.

We performed flexible molecular docking simulations (37, 38) between each of the nine GSE polyphenolic compounds and HuNoV VP1 proteins to discern stable docking conformations, across the four domains of VP1 (S, S-P1 hinge, P1, and P2 domain; **Supplementary Fig. 2**) to identify where each of the polyphenols tended to bind in order of domain preference. For these simulations, we used the HuNoV icosahedral asymmetric unit (PDB ID: 1IHM) that comprised three VP1 proteins because the complete HuNoV capsid consists of 180 identical icosahedral asymmetric units. Residue-level domain definitions have been described by Campillay-Véliz et al. (39). Flexible molecular docking simulations are shown in **Fig. 3**. The expected binding strength of each polyphenol (in increasing order of size/molecular weight) with HuNoV capsid protein has been reported where the error bars indicate variance across top 15 polyphenol binding poses within the same capsid groove. The scale on the right indicates fractional capsid-binding affinity of a given polyphenol with respect to the positive control – BSA. Larger polyphenols tend to show increased affinity towards both the positive control and target capsid of HuNoV. Analysis of binding affinities from docking experiments reveal greater electrostatic stabilization of the larger conjugated electron systems within the bigger polyphenols by the polar grooves of HuNoV capsid and bovine serum albumin alike. Data demonstrated that each of the nine polyphenolic compounds examined from GSE likely bound to different residues or regions of the HuNoV capsid protein. In addition, these polyphenols appeared to bind to a location at the dimeric interface of the trimeric capsid protein. The scores obtained from the simulation reflect the enthalpic contribution of binding (kcal/mol) between the polyphenols and the capsid (40). Fifteen independent trajectories of docking were performed using the docking protocol from OptMAVEn-2.0 suite (38) and the Rosetta energy function (41) was used to score the docked poses. The expected binding score was reported to be the modal value (i.e., most probable score) across the fifteen recorded values per complex. Since the polyphenols tend to show a conjugated electron system, larger polyphenols showed better electrostatic stabilization by the polar binding groove offered by the capsid proteins. Consequently, flavan-3-ol dimers showed stronger binding affinities to the capsid protein than smaller groups of polyphenols (**Fig. 3**). The polyphenol-capsid interaction profiles were composed primarily of hydrogen bonds along with hydrophobic interactions, accounting for the efficient capsid capture seen in experiments (**Fig. 4**). For example, the smallest polyphenol protocatechuate establishes three hydrogen bonded and three hydrophobic contacts while procyanidin (the largest polyphenolic compound) shows up to ten hydrogen bonded contacts yet maintaining the same number of hydrophobic interactions with the capsid (**Supplementary Fig. 3**). In addition to being concordant with a previous experimental study showing that polyphenols’ binding affinity to proteins (e.g., BSA and human salivary α-amylase) increased with their molecular weights (42), we also provide key biophysical insights for the same in this work (**Fig. 4**). Binding affinities for respective polyphenols with the HuNoV capsid were near-equal to that of bovine serum albumin, our positive control (**Fig. 3**). The bovine serum albumin is known as the major protein in FBS (43), with reported quenching activity towards polyphenolic compounds. Thus, our results indicated that the GSE polyphenols tend to promote capture of viral capsids with high efficiency.

**Fig. 3.**
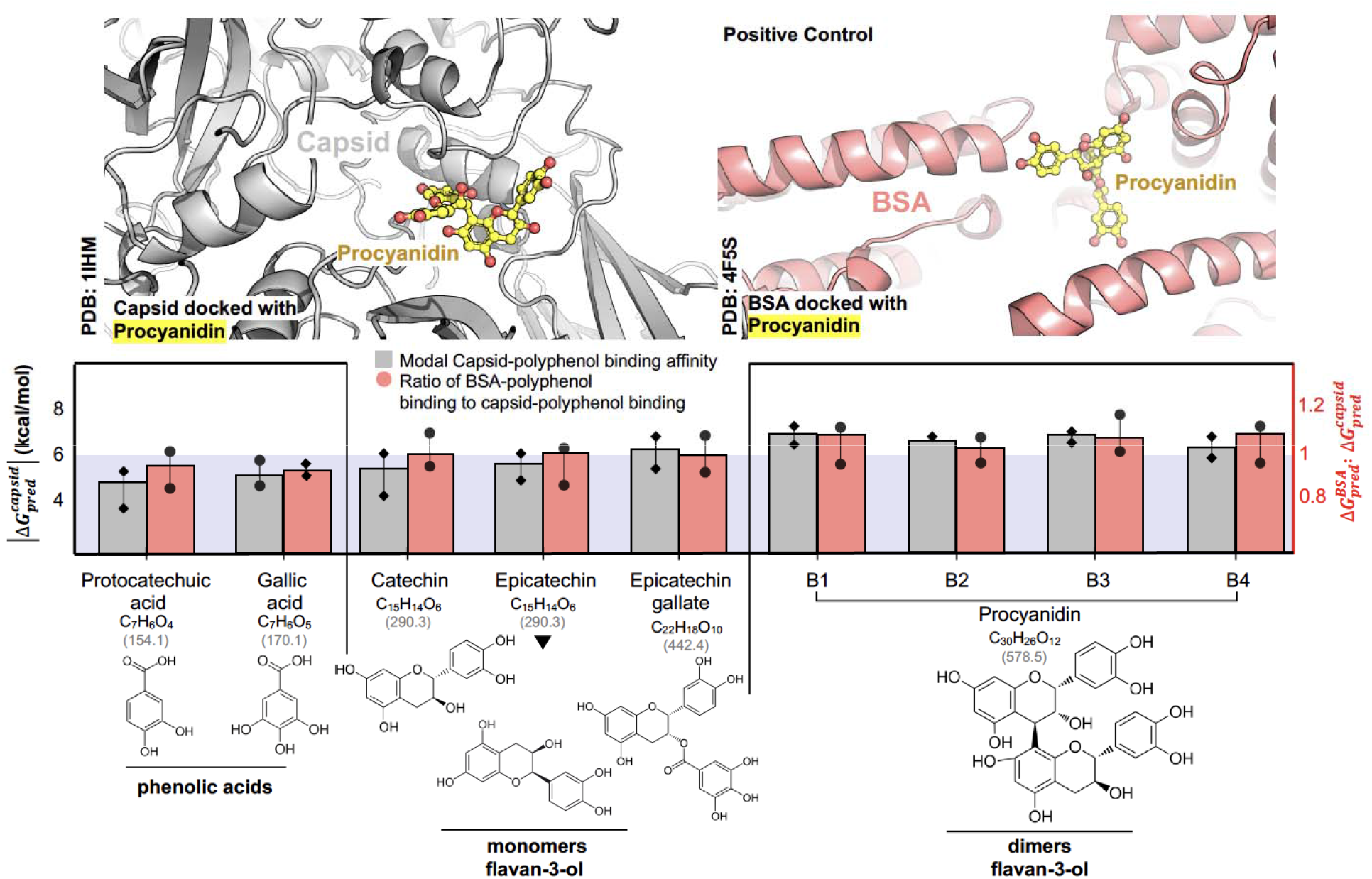
Summary of molecular docking analysis between HuNoV capsid and GSE selected polyphenols found in GSE. BSA, which is known to bind strongly with polyphenols (14), wa used as a positive control. The expected binding strength of each polyphenol (in increasing order of size/molecular weight) with HuNoV capsid protein (PDB id: 1IHM) has been reported where the error bars indicate variance across top 15 polyphenol binding poses within the same capsid groove. The scale on the right indicates fractional capsid-binding affinity of a given polyphenol with respect to the positive control – BSA.

**Fig. 4.**
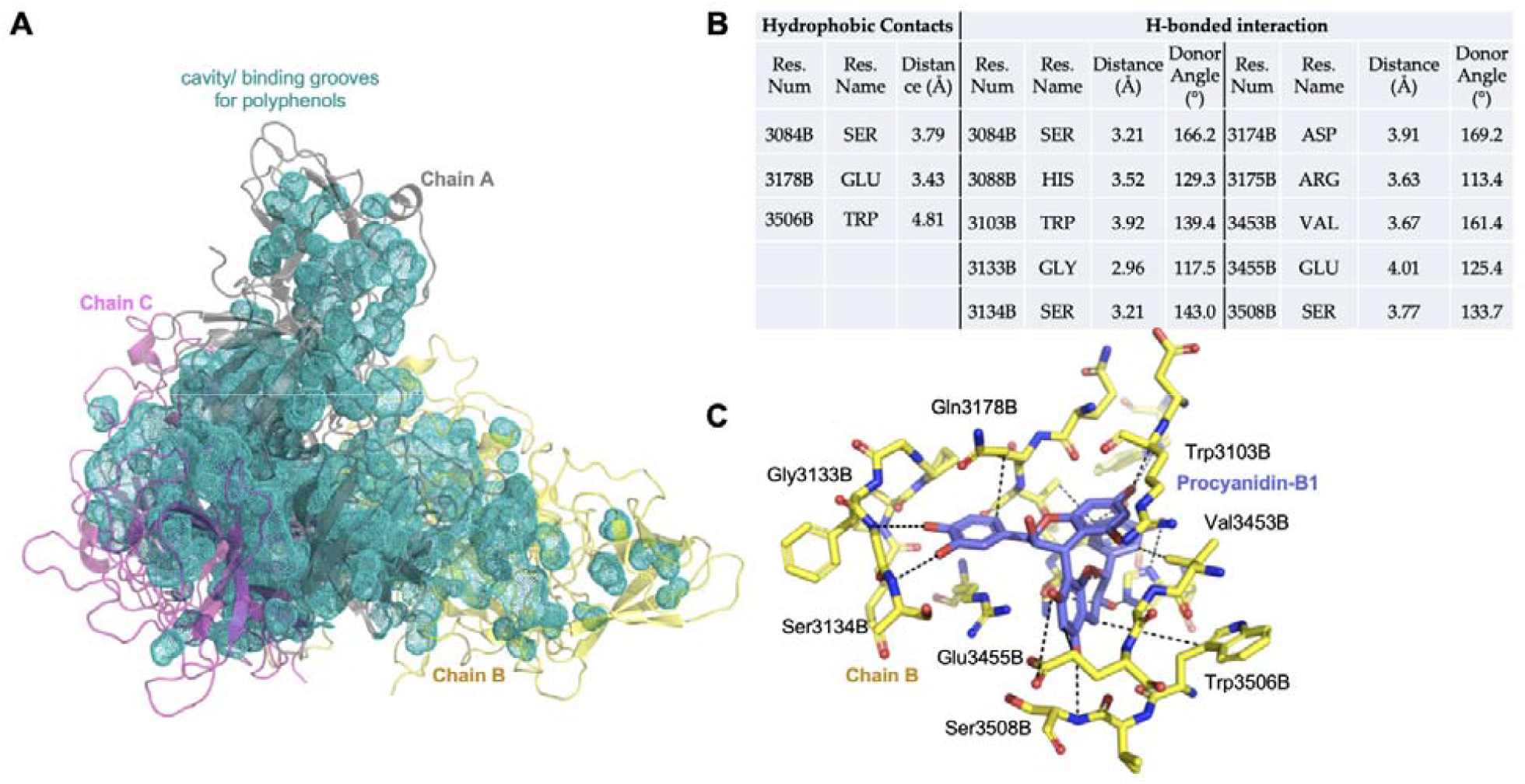
Larger polyphenols show greater electrostatic stabilization with the viral capsid. **(A)** HuNoV capsid shows a trigonal-symmetric polyphenol accessible groove (indicated in teal), which includes three intra-chain pockets, and three inter-chain binding crevices. Smaller polyphenols can access the inter-chain crevices, while the bigger ones remain localized to the more solvent-exposed intra-chain pockets. **(B)** Residue level interactions of Procyanidin-B1 with the HuNoV capsid are listed. **(C)** A graphical overview of the exact docked pose of Procyanidin-B1 in the capsid groove with most of the interactions is illustrated.

It is noteworthy that as the size of the polyphenol increases, it tends to bind to a less-buried, more solvent exposed pocket (owing to steric clashes) (**Fig. 5**) yet binding to the interface of two chains of the trimeric capsid (located around the S-P1 hinge domain; **Supplementary Fig. 2**). Only the largest Procyanidin (B1-B4) polyphenols cannot access the inter-chain interface pockets and are primarily surface binders with most interactions limited to a single chain (**Fig. 5**). We thereby hypothesize that larger polyphenols tend to be surface binders but stronger capsid binders, while smaller ones preferably bind to inter-chain interfaces of the viral capsid. In terms of domains bound, our docking simulations, in concordance with the experimental data, show that most polyphenols bind to the S or S-P1 hinge domains - while trace binding is observed in the P1 domain and P2 domain is mostly unbound.

**Fig. 5.**
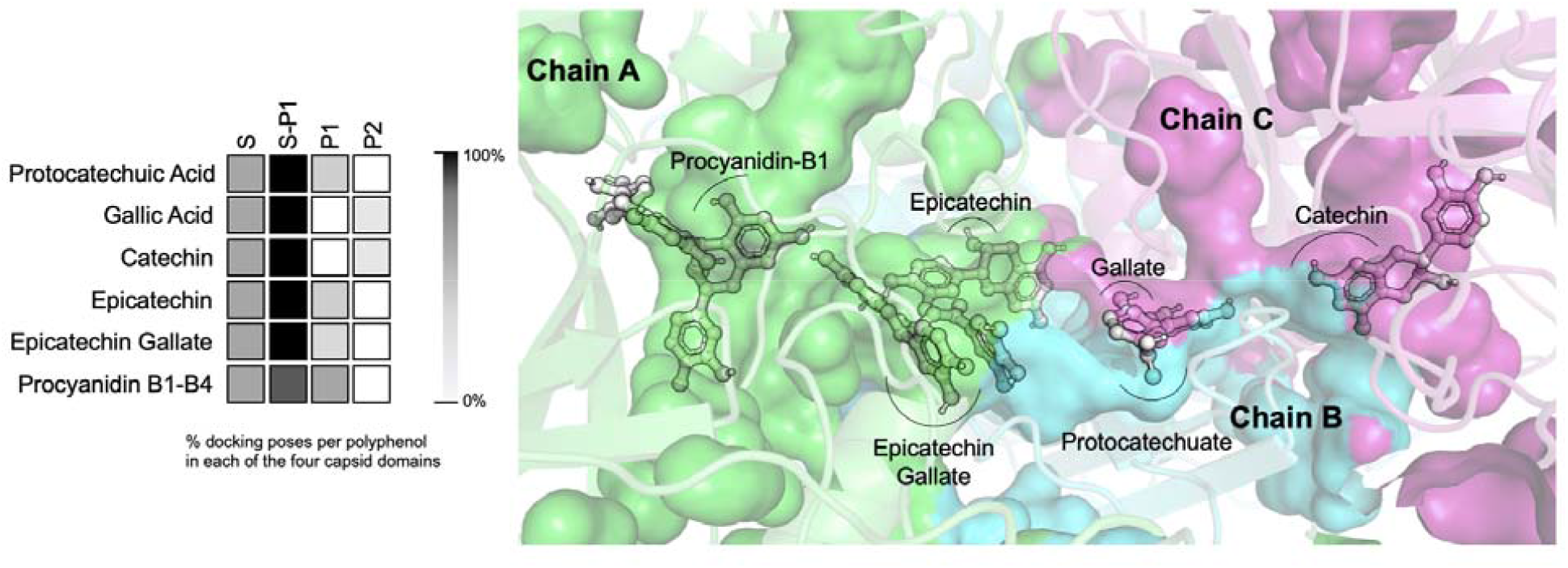
All polyphenol binding is primarily localized around the S and S-P1 hinge domains of the HuNoV capsid - while the P2 domain remains mostly unbound. Smaller polyphenols can access more buried inter-chain pockets of the HuNoV capsid, while larger Procyanidins bind to surface-exposed pockets with a single chain. Pockets guarded by each chain have been indicated by a different color (green – A, cyan – B, magenta – C). Smaller polyphenols – protocatechuate, gallate, and catechin bind best at the interface of chains B and C. Slightly larger epicatechin and epicatechin gallate bind best at the interface of chain A and B, while larger procyanidins (all B1 through B4) bind best to a solvent-exposed cavity within chain A only.

We extended the molecular docking simulations to other enteric viruses such as feline calicivirus (FCV-F9), murine norovirus (MNV-1), and hepatitis A virus (HAV) that were experimentally studied in previous works (14, 18, 19). Although the three sets of experiments were performed in different conditions such as different virus media (water or produce) and different pH, they presented a similar trend that FCV-F9 is more susceptible to the polyphenols in GSE than MNV-1 and HAV. Our molecular docking simulation agreed with the experimental data with these enteric viruses. The binding energy of the polyphenols to FCV-F9 capsid protein were stronger than those of MNV-1 and HAV (P<0.05) (**Fig. 6**). Note that the binding energy alone will not be sufficient information to evaluate the antiviral effect of the polyphenols because the binding energy does not necessarily indicate the following reactions such as viruses being aggregated and losing infectivity.

**Fig. 6.**
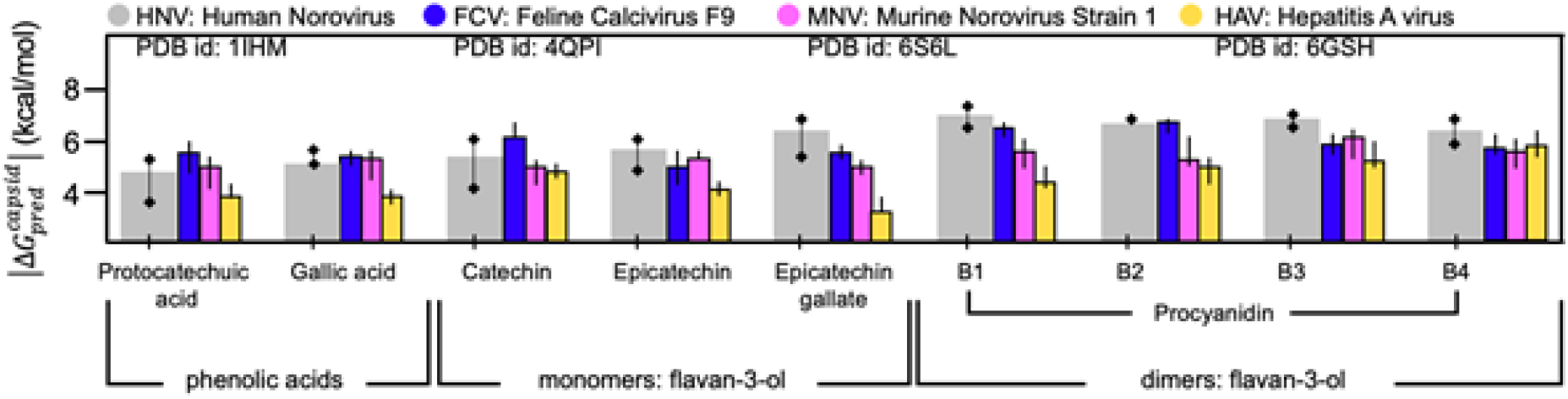
Expected binding energies (absolute modal values) between each polyphenolic compound and the four viral capsids (HuNoV: human norovirus, FCV: feline calicivirus F9, MNV: murine norovirus strain 1, and HAV: hepatitis A virus) were computed using the Rosetta binding energy function from top 15 docking poses per complex. The error bars indicate variance from the 15 docked poses for each complex. The reported energy scores were compared by a paired sample t-test indicating a statistically significant higher polyphenol binding activity by HuNoV and FCV in contrast to MNV (P<0.05) and HAV (p<0.001) capsids.

## Discussion

Anti-bacterial, anti-biofilm, and anti-fungal effects of plant-derived polyphenols are well-known. In this study, we systematically evaluated antiviral efficacy of the polyphenols in GSE to TV, a surrogate virus for HuNoV, and the antiviral mechanisms. Data shown here suggest that GSE can inactivate TV and perhaps other enteric viruses. In addition, data show that the polyphenols in GSE aggregate the virus particles, an event that is likely responsible for TV inactivation. Indeed, this concept that TV adsorbs polyphenols was also supported by the binding energy calculated by molecular docking analysis. Our current model is that TV aggregation prevents proper virus-host interactions or virus entry into host cells. There is at least one other report of aggregation of herpes simplex virus, prevents virus entry into cells while allowing virus attachment (44).

It should be noted that the TV concentrations used in this study are likely much higher than that found in contaminated produce or water. For example, rotavirus concentration in river water, treated wastewater, and untreated wastewater are reported to be 10^−3.0^, 10^−2.2^, and 10^−1.3^ FFU/mL respectively (45, 46). Given that GSE was efficacious using these artificially high TV concentrations used in the laboratory, it is reasonable to expect that GSE would be effective as an anti-viral for contaminated food or produce. Also, Tulane virus is more resistant to chemical disinfectants than the other viruses in the *Caliciviridae* family (47). Thus, if GSE were an effective disinfectant of Tulane virus then it is likely GSE would be efficacious against other caliciviruses, and this is an avenue of future research. GSE is also attractive because it is safe for consumption by humans. Thus, GSE or plant-derived polyphenols could be advantageous over the chemical disinfectants for direct addition to end-products that people ingest such as drinking water and foods.

There are still several important questions to answer before polyphenols can be widely used to inactivate the water-borne and food-borne pathogens. Water contains other organic matter which could outcompete viruses for binding to polyphenolic compounds. For example, Joshi et al. (14) showed the antiviral efficacy decreased when milk (which has high proteins) was added to an inactivation reaction. Thus, further studies need to be performed to understand how effective GSE is in water that has different properties. Other water properties (e.g., pH, ionic strength, and temperature) are important for adsorption reactions, and additional studies need to address how these properties may affect the efficacy of GSE. If GSE or individual polyphenols were to be used for foods (e.g., apples) then we must also understand how the surface properties of foods (e.g., wax contents of produce surface) may affect the efficacy of GSE (27).

## Materials and Methods

### Commercially available grape seed extract and its plant-derived polyphenols

We purchased a commercial grape seed extract (GSE) solution for this research (7832, Natures plus, USA). The total polyphenol (TP) concentration in each GSE sample was quantified by the colorimetric method using Folin-Ciocalteu reagent. A mixture of 100 μL GSE sample and 500 μL of 10% Folin-Ciocalteu phenol reagent (Sigma-Aldrich, MO, USA) was prepared in a 10 mm path length polymethyl methacrylate (PMMA) cuvette (BrandTech Scientific, CT, USA). Within 3 to 8 min of mixture preparation, 400 μL of 7.5% sodium carbonate solution was added to the mixture and homogenized by pipetting. After incubating at room temperature for 60 minutes, UV absorbance at 765 nm was measured by a spectrophotometer (UV-2450, Shimadzu, Japan). Distilled water was used as a reference solution. The same procedure was conducted with gallic acid solutions with different concentrations prepared by diluting gallic acid monohydrate (Sigma-Aldrich, MO, USA) in distilled water. The TP concentrations of the gallic acid solutions were used for a calibration curve (**Supplementary Fig. 4**) which determined TP concentration of the GSE samples. GSE with TP concentrations ranging from 84 to 1694 μg/mL was prepared by diluting the initial GSE with 1X PBS and used for inactivation experiments.

The polyphenol composition of GSE was analyzed by liquid chromatography-mass spectrometry (Q Exactive™ Plus Hybrid Quadrupole-Orbitrap™ Mass Spectrometer, Thermo scientific, USA), and numerous peaks were discovered by untargeted comprehensive metabolite profiling (**Supplementary Table 2**). Although the LC/MS analysis could not quantify each polyphenol and differentiate polyphenolic isomers with the same m/z values, such as catechin/epicatechin at 291 of m/z ratio for [M+H]^+^ and B type procyanidins at 579 of m/z ratio for [M+H]^+^, the major peaks agreed with the primary polyphenolic compounds in GSE that were confirmed by other studies (35, 36). With the mass spectrometry and the references characterizing polyphenols in GSE, we chose nine target polyphenolic components, which include three monomers flavan-3-ols (catechin, epicatechin, and epicatechin gallate), four dimers flavan-3-ols (Procyanidin B1, Procyanidin B2, Procyanidin B3, and Procyanidin B4), and two phenolic acid (gallic acid and protocatechuic acid). These nine polyphenolic compounds were further studied through molecular docking simulations.

### Tulane virus propagation

Tulane virus (TV) was used as a surrogate for human norovirus (HuNoV). Both TV and HuNoV are members of *Caliciviridae* and share structural similarities. TV and HuNoV have a positive-sense, single-stranded genomic RNA of 6.7 kb and 7.5 kb, respectively. About 23-26% of the genome sequences are identical in both viruses (20). The capsid of TV has T=3 icosahedral symmetry and is 40 nm in diameter. The capsid comprises 180 copies of major capsid protein (VP1) or 90 dimers of A/B or C/C. The VP1 is also sub-divided into S, P1, and P2 domains. When the VP1 proteins are assembled, the S domains comprise the bottom surface of the capsid while P1 and P2 domains protrude out of the bottom surface, which is responsible for binding to receptor proteins (**Supplementary Fig. 1**). TV recognizes Histo-blood group antigens (21) and sialic acids (22) as cellular receptors similar to HuNoVs. Besides, TV is considered more resistant to disinfectants than the other viruses in *Caliciviridae* (47). Therefore, TV has been widely used in virus inactivation experiments as a surrogate for HuNoVs (7, 23–27).

TV was a gift from Cincinnati Children’s Hospital Medical Center (22) and the TV genome was sequenced as quality control. The genome sequence was 100% identical to those of a wild-type TV strain in Genbank (Access number: EU391643) (7). TV was propagated in MA104 cells, which was purchased from ATCC (CRL-2378.1), and grown in complete culture medium (i.e., 1X minimum essential medium (MEM; Thermo Fisher Scientific, USA), 10% fetal bovine serum (FBS; Thermo Fisher Scientific, USA), 1X antibiotic-antimycotic (Thermo Fisher Scientific, USA), 17 mM of NaHCO_3_, 10 mM of HEPES, and 1 mM of sodium pyruvate (Thermo Fisher Scientific, USA). When greater than 90% of MA014 cellular monolayers showed cytopathic effects (about 2 days after the inoculation), cells were harvested and collected by centrifugation, and TV was released from host cells by three cycles of freeze and thaw. Virus was separated from cellular debris by centrifugation at 2000 rpm (556 g) for 10 min (Sorvall Legend RT Plus, Thermo Fisher Scientific, USA). The supernatant was treated by filtration with a 0.22 μm bottle top filter (Milliporesigma, USA) to remove additional cellular debris. The virus-containing filtrate was then further purified using an ultracentrifuge (Optima XPN-90 Ultracentrifuge, Beckman Coulter, USA) with a cycle of 1,000 rpm (116 g) for 5 min followed by 36,000 rpm (150700 g) at 4°C for 3 hours. The virus pellet was resuspended in 1X PBS, aliquoted, and stored at −80LJ before use.

### Tulane virus inactivation experiments using GSE

Virus inactivation experiments were initiated by adding 250 μL of TV solution containing 2.5 ×10^5^ PFU TV to 250 μL of GSE-containing solution, in which GSE concentrations ranged from 84 to 1694 μg/mL. After the incubation times indicated in the manuscript (i.e., 10 to 120 seconds), 70 μL of the mixture was added to 70 μL of FBS to quench the polyphenolic activity(14). Thus, the volumetric ratio of TV, GSE, and FBS in the final solution was 1:1:2. The FBS quenching activity was confirmed in **Supplementary Fig. 5**. Negative controls were prepared for every virus inactivation experiment. In this case, the negative controls were prepared by mixing TV, GSE, and FBS in the ratio of 1:1:2, but in a different mixing order. Specifically, 35 μL of GSE was mixed with 70 μL of FBS to quench the polyphenolic activity followed by adding 35 μL of TV to the mixture (**Fig. 1A**). The final mixture of TV, GSE, and FBS was used for further analysis.

### Plaque assays

The MA104 cell line was grown in 175 cm^2^ flasks (Thermo Fisher Scientific, USA) with the complete culture medium. Cells were seeded into 6-well plates (CC7682-7506, USA Scientific, USA) resulting in cellular monolayers with more than 90% confluency. Cell culture supernatants were aspirated using a vacuum-connected pasteur pipette. 100 μL of TV samples were serially diluted by 10-fold dilutions. Cellular monolayers were incubated with each of the serially-diluted viruses for 1 hour at 37□ with 5% CO_2_ to facilitate virus attachment to the MA104 cells. Viruses were aspirated from cellular monolayers and 2 mL of overlay solution containing 1X MEM, 1% agarose, 7.5% sodium bicarbonate, 15 mM HEPES, and 1X antibiotic-antimycotic was added to each well. The overlay solution was solidified at 4□ for 10 minutes. Plates were incubated for 2 days at 37□ with 5% CO_2_ to allow infectious viruses to form plaques. Next, 2 mL of 10% formaldehyde (VWR, USA) in 1X PBS was added to each well and incubated at room temperature for 1 hour to fix cells. The agarose and the formaldehyde were removed and replaced with a 0.05% crystal violet dye solution. The solution was washed away after 10 minutes and the number of plaque-forming units (PFU) was counted on a lightbox (ULB-100, Scienceware). The detection limit of the plaque assay was one plaque on the least diluted sample (i.e., 10-fold dilution), which was equivalent to 10^1.1^ PFU/mL.

### Models for virus inactivation kinetics

TV inactivation kinetics were interpreted by a first-order reaction of Chick’s law (Eqs. 1-3) and pseudo-second-order model (Eqs. 4-6). Chick’s law assumes the activity (i.e., concentration) of available disinfectant remains constant during the reaction. Chick’s law has been widely used to describe virus inactivation kinetics by free chlorine, peracetic acid, monochloramine, heat, ozone, and UV(7, 48–51).

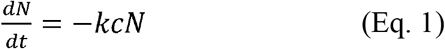

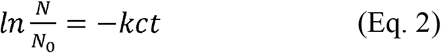

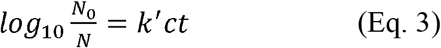

Where *N* referred to viral infectivity (PFU/mL), *c* indicated the TP concentration (μg/mL), *t* was treatment time (s), *k*^*′*^ meant an inactivation rate constant from Chick’s law (L/μg⋅s).

The pseudo-second-order model has been widely used to describe reactions where chemisorption between adsorbent and adsorbate is the rate-determining step (28). In this study, TV and TP were considered as adsorbate and adsorbent, respectively.

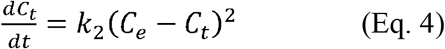

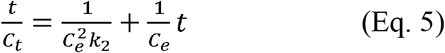

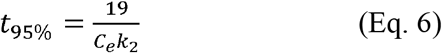

Where t is the reaction time (s), k_2_ is the rate constant from the pseudo-second-order model (mL/PFU·s), C_t_ and C_e_ are inactivated TV concentration (PFU/mL) at time t and equilibrium state, respectively. Also, t_95%_ is a reaction time at which C_t_ reaches 95% of C_e_.

### Virus particle size

The particle size distribution of TV was measured by a light scattering analyzer (DelsaMax Pro, Beckman Coulter). TV, GSE, and FBS either alone or in combination were prepared in 200 μL aliquots, placed in a PMMA cuvette (BrandTech Scientific, USA). Particle size was measured 20 times for each sample, and the averaged % intensity was presented with the particle diameter. According to the manufacturer, particle size analysis is reliable within a range from 0.4 to 10,000 nm, and all our measurements were in this detection range.

### Virus aggregation assay

An assay was developed to quantify TV virions that are less than 100 μm in diameter. After the TV inactivation experiments were conducted, the mixture containing viruses, GSE, and FBS was filtered with a 0.1 μm syringe filter (Sartorius, Germany). The number of virus particles in the initial mixture and the filtrate was quantified by one-step RT-qPCR. RNA was extracted from samples using Viral RNA Mini Kit (QIAGEN, Germany) in a final 60 μL volume. RT-qPCR samples were prepared in 96-well plates (4306737, Applied Biosystems, USA) by mixing 5 μL of 2 × iTaq universal SYBR green reaction mix, 0.125 μL of iScript reverse transcriptase from the iTaq universal SYBR green reaction mix (Bio-Rad Laboratories, USA), 3 μL of the RNA, 0.3 μL of 10 μM TV-NSP1-qPCR-F primer, 0.3 μL of 10 μM TV-NSP1-qPCR-R primer, and 1.275 μL of nuclease-free water. The one-step RT-qPCR was run using a qPCR system (QuantStudio 3, Thermo Fisher Scientific, USA) with the following thermocycle: 10 minutes at 50□ and 1 minute at 90□ followed by 40 cycles of 30 seconds at 60□ and 1 minute at 90□. Detailed information for primers and synthetic DNA controls is summarized in **Supplementary Table 3**. We obtained a calibration curve by plotting the viral infectivity (Log_10_ PFU/mL) on the x-axis and the outcome of the one step RT-qPCR assay (Log_10_ gc/mL) on the y-axis for the same viral solution (**Supplementary Fig. 6**). The calibration curve was used to determine the reliable range of the binding assay. The calibration curve for the binding assay was linear (a slope=0.95 and R^2^=1.00) between 10^3^ and 10^7^ PFU/mL. The PCR standard curves were obtained with 10-fold serial dilutions of the synthetic DNA oligonucleotide (Integrated DNA technologies, USA) and the PCR efficiency for this one-step RT-qPCR ranged from 85 to 95% (R^2^>0.99). Inhibitory effect of GSE was also tested (**Supplementary Fig. 7**). We found 100-fold dilution can reduce the inhibitory effect of GSE to an insignificant level, so the samples were diluted in molecular grade water by 100-fold before the RT-qPCR analysis. Information for RT-qPCR was summarized following the MIQE guidelines (52) in **Supplementary Table 4**.

### Molecular docking simulation for polyphenols and capsid proteins interaction

We selected nine polyphenolic compounds that are present in GSE, including two phenolic acids, three monomer flavan-3-ols, and four dimer flavan-3-ols (36). The structural information on the polyphenols were obtained from PUBChem (https://pubchem.ncbi.nlm.nih.gov/). Although we conducted inactivation experiments with Tulane virus, we obtained HuNoV capsid protein instead of TV from the Protein Data Bank (PDB, https://www.rcsb.org/) to study interactions with the polyphenols because the database of PDB does not provide TV capsid information. Cryo-EM structure analysis showed that the TV capsid structure closely resembles that of HuNoV(31).

We downloaded the HuNoV icosahedral asymmetric unit (PDB ID: 1IHM) which is a basic building block for the HuNoV capsid. This icosahedral unit comprised three VP1 proteins (i.e., chain A, B, and C in PDB format). The complete HuNoV capsid consists of 180 identical icosahedral asymmetric units. We also used a deep-learning-based structure prediction tool - trRosetta(53), to predict the 3D-structure of the TV capsid (**Supplementary File 1**) from its sequence and a multiple-sequence alignment of related sequences. We studied molecular docking of the polyphenols to HuNoV and to other different enteric viruses including hepatitis A virus (PDB: 4QPI), murine norovirus-1 (PDB: 6S6L), and feline calicivirus-F9 (PDB: 6GSH). Bovine serum albumin (BSA) is a major component for FBS, and BSA showed a strong binding affinity to the polyphenols as confirmed by the quenching effect in the virus inactivation experiments. BSA (PDB: 4F5S) was used as a positive control for molecular docking with the polyphenolic compounds. Next, we used the target proteins (capsids or BSA) and flexible conformations of all the aforementioned nine polyphenol ligands to discern stable docking conformations, record binding affinity scores, and report across four capsid domains where each of the polyphenols tend to bind in order of domain preference. The flexible docking protocol is similar to the Z-Dock protocol(37) as implemented within OptMAVEn-2.0(38).

### Statistical analysis

All experiments were repeated three times with distinct virus samples (i.e., three biological replications), each of which was analyzed by three separate RT-qPCR measurements (i.e., three technical replications). Paired sample t-test was used to compare results of plaque assay and aggregation assay in **Fig. 2C** and binding energies of different viral species (HuNoV, FCV, MNV, and HAV) to polyphenolic compounds in **Fig. 6**. We confirmed that all data for statistical analysis satisfied assumptions of paired sample t-test (i.e., no outliers and normality). For example, all data for statistical analysis were between Q1-1.5IQR and Q3+1.5IQR range, which means there were no outliers. Also, differences between two data sets (e.g., plaque assay versus aggregation assay in **Fig. 2C** or HuNoV versus MNV or HAV in **Fig. 6**) were normally distributed (P>0.05).

## Acknowledgement

This study was funded by National Science Foundation (Award Number: 2023248). We thank Dr. Zhong (Lucas) Li at Metabolomics Lab, Roy J. Carver Biotechnology Center, University of Illinois at Urbana-Champaign for characterizing polyphenolic compounds in grape seed extract.

## References

1. Ahmed SM, Hall AJ, Robinson AE, Verhoef L, Premkumar P, Parashar UD, Koopmans M, Lopman BA. 2014. Global prevalence of norovirus in cases of gastroenteritis: A systematic review and meta-analysis. Lancet Infect Dis 14:725–730.

2. Scallan E, Hoekstra RM, Angulo FJ, Tauxe R V., Widdowson MA, Roy SL, Jones JL, Griffin PM. 2011. Foodborne illness acquired in the United States-Major pathogens. Emerg Infect Dis 17:7–15.

3. Burke RM, Mattison CP, Pindyck T, Dahl RM, Rudd J, Bi D, Curns AT, Parashar U, Hall AJ. 2020. Burden of Norovirus in the United States, as Estimated Based on Administrative Data: Updates for Medically Attended Illness and Mortality, 2001–2015. Clin Infect Dis https://doi.org/10.1093/cid/ciaa438.

4. Bitler EJ, Matthews JE, Dickey BW, Eisenberg JNS, Leon JS. 2013. Norovirus outbreaks: a systematic review of commonly implicated transmission routes and vehicles. Epidemiol Infect 141:1563–1571.

5. Fraisse A, Temmam S, Deboosere N, Guillier L, Delobel A, Maris P, Vialette M, Morin T, Perelle S. 2011. Comparison of chlorine and peroxyacetic-based disinfectant to inactivate Feline calicivirus, Murine norovirus and Hepatitis A virus on lettuce. Int J Food Microbiol 151:98–104.

6. Girard M, Ngazoa S, Mattison K, Jean J. 2010. Attachment of Noroviruses to Stainless Steel and Their Inactivation, Using Household Disinfectants. J Food Prot 73:400–404.

7. Fuzawa M, Araud E, Li J, Shisler JL, Nguyen TH. 2019. Free Chlorine Disinfection Mechanisms of Rotaviruses and Human Norovirus Surrogate Tulane Virus Attached to Fresh Produce Surfaces. Environ Sci Technol 53:11999–12006.

8. Chen X, Hung YC. 2016. Predicting chlorine demand of fresh and fresh-cut produce based on produce wash water properties. Postharvest Biol Technol 120:10–15.

9. Parish ME, Beuchat LR, Suslow T V., Harris LJ, Garrett EH, Farber JN, Busta FF. 2003. Methods to reduce/ eliminate pathogens from fresh and fresh-cut produce. Compr Rev Food Sci Food Saf 2:161–173.

10. Soto Beltran M, Jimenez Edeza M, Viera C, Martinez CI, Chaidez C. 2013. Sanitizing alternatives for Escherichia coli and Salmonella typhimurium on bell peppers at household kitchens. Int J Environ Health Res 23:331–341.

11. Matulonga B, Rava M, Siroux V, Bernard A, Dumas O, Pin I, Zock JP, Nadif R, Leynaert B, Le Moual N. 2016. Women using bleach for home cleaning are at increased risk of non-allergic asthma. Respir Med 117:264–271.

12. Cleveland J, Montville TJ, Nes IF, Chikindas ML. 2001. Bacteriocins: Safe, natural antimicrobials for food preservation. Int J Food Microbiol. Elsevier.

13. Devlieghere F, Vermeiren L, Debevere J. 2004. New preservation technologies: Possibilities and limitations, p. 273–285. In International Dairy Journal. Elsevier.

14. Joshi SS, Su X, D’Souza DH. 2015. Antiviral effects of grape seed extract against feline calicivirus, murine norovirus, and hepatitis A virus in model food systems and under gastric conditions. Food Microbiol 52:1–10.

15. Market analysis report. 2019. Polyphenols Market Size, Share & Trends Analysis Report By Product (Grape Seed, Green Tea, Cocoa), By Application (Beverages, Food, Feed, Dietary Supplements, Cosmetics), And Segment Forecasts, 2019 –2025.

16. Memar MY, Adibkia K, Farajnia S, Kafil HS, Yekani M, Alizadeh N, Ghotaslou R. 2019. The grape seed extract: A natural antimicrobial agent against different pathogens. Rev Med Microbiol 30:173–182.

17. D’Souza DH. 2014. Phytocompounds for the control of human enteric viruses. Curr Opin Virol 4:44–49.

18. Su X, D’Souza DH. 2011. Grape seed extract for control of human enteric viruses. Appl Environ Microbiol 77:3982–3987.

19. Su X, D’Souza DH. 2013. Grape seed extract for foodborne virus reduction on produce. Food Microbiol 34:1–6.

20. Farkas T, Sestak K, Wei C, Jiang X. 2008. Characterization of a Rhesus Monkey Calicivirus Representing a New Genus of Caliciviridae. J Virol 82:5408–5416.

21. Zhang D, Huang P, Zou L, Lowary TL, Tan M, Jiang X. 2015. Tulane Virus Recognizes the A Type 3 and B Histo-Blood Group Antigens. J Virol 89:1419–1427.

22. Tan M, Wei C, Huang P, Fan Q, Quigley C, Xia M, Fang H, Zhang X, Zhong W, Klassen JS, Jiang X. 2015. Tulane virus recognizes sialic acids as cellular receptors. Sci Rep 5:1–14.

23. Bridges DF, Breard A, Lacombe A, Valentine DC, Tadepalli S, Wu VCH. 2017. Inhibition of Tulane Virus replication via exposure to lowbush blueberry (Vaccinium angustifolium) fractional components. J Berry Res 7:281–289.

24. Li X, Huang R, Chen H. 2017. Evaluation of Assays to Quantify Infectious Human Norovirus for Heat and High-Pressure Inactivation Studies Using Tulane Virus. Food Environ Virol 9:314–325.

25. Ailavadi S, Davidson PM, Morgan MT, D’Souza DH. 2019. Thermal Inactivation Kinetics of Tulane Virus in Cell-Culture Medium and Spinach. J Food Sci 84:557–563.

26. Araud E, Fuzawa M, Shisler JL, Li J, Nguyen TH. 2020. UV inactivation of rotavirus and tulane virus targets different components of the virions. Appl Environ Microbiol 86:1–12.

27. Fuzawa M, Bai H, Shisler JL, Nguyen TH. 2020. The basis of peracetic acid (PAA) inactivation mechanisms for rotavirus and Tulane virus under conditions relevant for vegetable sanitation. Appl Environ Microbiol 1–47.

28. Ho YS, McKay G. 1999. Pseudo-second order model for sorption processes. Process Biochem 34:451–465.

29. Ho YS, Ng JCY, McKay G. 2001. Removal of lead(II) from effluents by sorption on peat using second-order kinetics. Sep Sci Technol 36:241–261.

30. Ho YS, Ofomaja AE. 2006. Pseudo-second-order model for lead ion sorption from aqueous solutions onto palm kernel fiber. J Hazard Mater 129:137–142.

31. Yu G, Zhang D, Guo F, Tan M, Jiang X, Jiang W. 2013. Cryo-EM Structure of a Novel Calicivirus, Tulane Virus. PLoS One 8:59817.

32. Pianet I, André Y, Ducasse M-A, Tarascou I, Lartigue J-C, Pinaud N, Fouquet E, Dufourc EJ, Laguerre M. 2008. Modeling Procyanidin Self-Association Processes and Understanding Their Micellar Organization: A Study by Diffusion NMR and Molecular Mechanics. Langmuir 24:11027–11035.

33. Islam B, Sharma C, Adem A, Aburawi E, Ojha S. 2015. Insight into the mechanism of polyphenols on the activity of HMGR by molecular docking. Drug Des Devel Ther 9:4943–4951.

34. Srinivasan E, Rajasekaran R. 2016. Computational investigation of curcumin, a natural polyphenol that inhibits the destabilization and the aggregation of human SOD1 mutant (Ala4Val). RSC Adv 6:102744–102753.

35. Chedea VS, Echim C, Braicu C, Andjelkovic M, Verhe R, Socaciu C. 2011. Composition in polyphenols and stability of the aqueous grape seed extract from the romanian variety “merlot recas.” J Food Biochem 35:92–108.

36. Rodríguez Montealegre R, Romero Peces R, Chacón Vozmediano JL, Martínez Gascueña J, García Romero E. 2006. Phenolic compounds in skins and seeds of ten grape Vitis vinifera varieties grown in a warm climate. J Food Compos Anal 19:687–693.

37. Pierce BG, Hourai Y, Weng Z. 2011. Accelerating protein docking in ZDOCK using an advanced 3D convolution library. PLoS One 6.

38. Chowdhury R, Allan MF, Maranas CD. 2018. OptMAVEn-2.0: De novo Design of Variable Antibody Regions Against Targeted Antigen Epitopes. Antibodies 7:23.

39. Campillay-Véliz CP, Carvajal JJ, Avellaneda AM, Escobar D, Covián C, Kalergis AM, Lay MK. 2020. Human Norovirus Proteins: Implications in the Replicative Cycle, Pathogenesis, and the Host Immune Response. Front Immunol. Frontiers Media S.A.

40. Cosconati S, Forli S, Perryman AL, Harris R, Goodsell DS, Olson AJ. 2010. Virtual screening with AutoDock: Theory and practice. Expert Opin Drug Discov. NIH Public Access.

41. Chaudhury S, Lyskov S, Gray JJ. 2010. PyRosetta: A script-based interface for implementing molecular modeling algorithms using Rosetta. Bioinformatics. Oxford Academic.

42. Soares S, Mateus N, De Freitas V. 2007. Interaction of different polyphenols with Bovine Serum Albumin (BSA) and Human Salivary α-Amylase (HSA) by fluorescence quenching. J Agric Food Chem 55:6726–6735.

43. Johnson M. 2012. Fetal Bovine Serum. Mater Methods 2.

44. Bultmann H, Busse JS, Brandt CR. 2001. Modified FGF4 Signal Peptide Inhibits Entry of Herpes Simplex Virus Type 1. J Virol 75:2634–2645.

45. Lodder WJ, De Roda Husman AM. 2005. Presence of noroviruses and other enteric viruses in sewage and surface waters in The Netherlands. Appl Environ Microbiol 71:1453–1461.

46. Rutjes SA, Lodder WJ, Van Leeuwen AD, De Roda Husman AM. 2009. Detection of infectious rotavirus in naturally contaminated source waters for drinking water production. J Appl Microbiol 107:97–105.

47. Cromeans T, Park GW, Costantini V, Lee D, Wang Q, Farkas T, Lee A, Vinjé J. 2014. Comprehensive comparison of cultivable norovirus surrogates in response to different inactivation and disinfection treatments. Appl Environ Microbiol 80:5743–5751.

48. Dunkin N, Weng S, Schwab KJ, McQuarrie J, Bell K, Jacangelo JG. 2017. Comparative Inactivation of Murine Norovirus and MS2 Bacteriophage by Peracetic Acid and Monochloramine in Municipal Secondary Wastewater Effluent. Environ Sci Technol 51:2972–2981.

49. Oguma K, Kita R, Sakai H, Murakami M, Takizawa S. 2013. Application of UV light emitting diodes to batch and flow-through water disinfection systems. Desalination 328:24–30.

50. Araud E, DiCaprio E, Ma Y, Lou F, Gao Y, Kingsley D, Hughes JH, Li J. 2016. Thermal inactivation of enteric viruses and bioaccumulation of enteric foodborne viruses in live oysters (Crassostrea virginica). Appl Environ Microbiol 82:2086–2099.

51. Dong S, Li J, Kim MH, Park SJ, Eden JG, Guest JS, Nguyen TH. 2017. Human health trade-offs in the disinfection of wastewater for landscape irrigation: Microplasma ozonation: Vs. chlorination. Environ Sci Water Res Technol 3:106–118.

52. Bustin SA, Benes V, Garson JA, Hellemans J, Huggett J, Kubista M, Mueller R, Nolan T, Pfaffl MW, Shipley GL, Vandesompele J, Wittwer CT. 2009. The MIQE guidelines: Minimum information for publication of quantitative real-time PCR experiments. Clin Chem 55:611–622.

53. Yang J, Anishchenko I, Park H, Peng Z, Ovchinnikov S, Baker D. 2020. Improved protein structure prediction using predicted interresidue orientations. Proc Natl Acad Sci U S A 117:1496–1503.

