## Supplemental Figures and Tables for "Inactivation mechanism and efficacy of grape seed extract for Human Norovirus surrogate"

Running title: Grape seed extract inactivates Tulane viruses

Chamteut Oh^1,*^, Ratul Chowdhury^2,*^, Laxmicharan Samineni^3^, Joanna L Shisler^4^, Manish Kumar^6^, and Thanh H. Nguyen^1,5^

^1^Department of Civil and Environmental Engineering, University of Illinois at Urbana-Champaign, USA

^2^Department of Chemical Engineering, The Pennsylvania State University, PA, USA

^3^Department of Chemical Engineering, The University of Texas at Austin, Austin, USA

^4^Department of Microbiology, University of Illinois at Urbana-Champaign, USA

^5^Institute of Genomic Biology, University of Illinois at Urbana-Champaign, USA

^6^Department of Civil, Architectural and Environmental Engineering, The University of Texas at Austin, Austin, USA

*^*^Equal contribution*

**Seven Supplementary Figures**

**Four Supplementary Tables**

**One Supplementary Text**

**Supplementary Table 1.** Kinetics parameters from inactivation experiments with different total polyphenol concentrations

| **GSE**  **(μg/mL)** | **Pseudo-second-order model** | | | | **Chick’s law** |
| --- | --- | --- | --- | --- | --- |
|  | **K_2_ (mL/PFU⋅s)** | **t_95%_ (s)** | **q_e_ (PFU/μg)** | **R^2^** | **R^2^** |
| 42 | 1.9$\times$10^-6^ | 1.2$\times$10^3^ | 8.6$\times$10^3^ | 0.25 | 0.07 |
| 169 | 5.2$\times$10^-5^ | 5.8$\times$10^1^ | 6.3$\times$10^3^ | 0.99 | 0.44 |
| 296 | 1.6$\times$10^-3^ | 1.8$\times$10^0^ | 6.8$\times$10^3^ | 1.00 | 0.54 |
| 424 | 2.4$\times$10^-2^ | 2.2$\times$10^-1^ | 3.6$\times$10^3^ | 1.00 | 0.34 |
| 551 | 7.5$\times$10^-2^ | 1.2$\times$10^-1^ | 2.1$\times$10^3^ | 1.00 | 0.44 |
| 678 | 1.6$\times$10^-1^ | 3.7$\times$10^-2^ | 3.2$\times$10^3^ | 1.00 | 0.43 |

**Supporting Text 1 (Validation experiment for aggregation assay)**

One concern for aggregation assay was that diluting GSE concentrations and vortexing virus solutions, which were main activities for plaque assay, could reverse the aggregation effects induced by GSE, resulting in virus segregation(1–4). If the GSE-induced virus aggregates are dispersed during plaque assay and become infectious virus particles, it potentially would confound the comparison between plaque assay and aggregation assay. Thus, we performed an experiment to examine the impact of vortexing and serial dilution on GSE-induced virion aggregates. For these experiments, TV was inactivated by 847 TP-μg/mL of GSE, which was the highest GSE concentration for the virus aggregation assay (**Fig. 1B**), and subsequently quenched by FBS. Next, samples were serially diluted in complete culture media and vortexed for 10 seconds to mimic processes for plaque assay. These serial dilutions were examined by one-step RT-qPCR to represent the total number of virions that were aggregated by GSE and then diluted and vortexed (Black symbols in **Supplementary Fig. 1**). The results are located on the 1:1 line in Supplementary Fig. 1, which means the virus aggregation alone did not affect the total virion numbers. Each serial dilution was then subjected to filtration, using filters with a 0.1 μm size pore. The filtrates were also quantified by one-step RT-qPCR to show the number of virions that passed through the 0.1 μm filter, and therefore were no longer aggregated, after the above-mentioned processes (Red symbols in **Supplementary Fig. 1**). Considering a single TV virion diameter of 40 nm(5), TVs that are aggregated with three or more virions will be excluded by the 0.1 μm pore size filter. Note that when intact TV was mixed with PBS, instead of the polyphenols, the number of virion aggregates larger than 0.1 μm accounted for less than 0.3-log_10_. Thus, the differences in those two values (i.e., gap between the black and red symbols in **Supplementary Fig. 1**) means the number of virion aggregates with a diameter of larger than 0.1 μm. If the virions were segregated back to single virus particles during the 10-fold serial dilution and the 10-second vortex, the number of virions that can pass through the 0.1 μm filter would have increased, and the number of virions after the filtration (red in **Supplementary Fig. 1**) would have gotten closer to that before the filtration (black in **Supplementary Fig. 1**). However, it did not happen over the three cycles of 10-fold dilution and the 10-second vortex. Therefore, the aggregated virions will not be segregated back to single virions during the plaque assay.

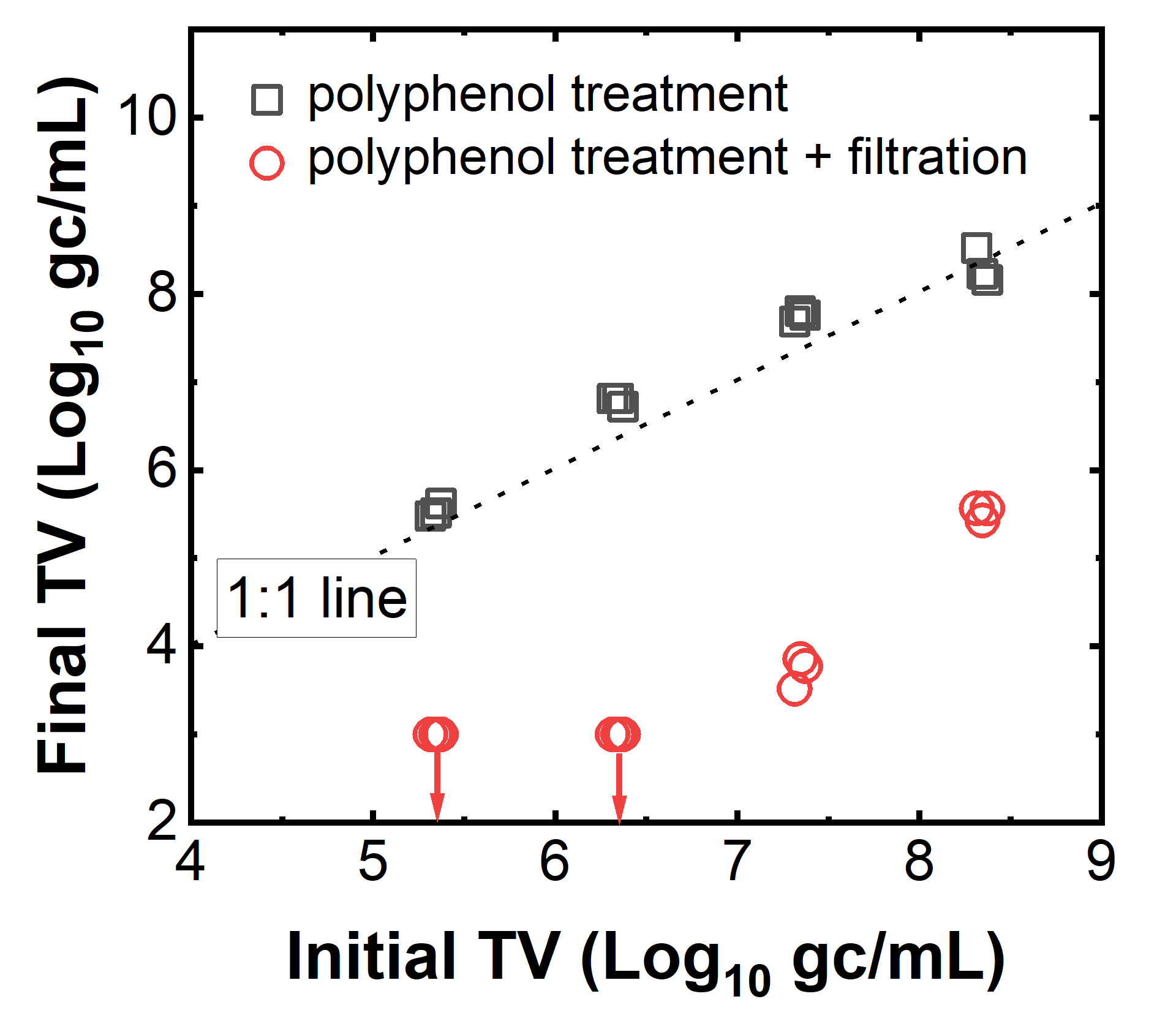

**Supplementary Fig. 1.** The impact of vortexing and dilution manipulations on virion aggregates. Black symbols indicate the number of virions that were aggregated by GSE, followed by being serially diluted and vortexed. Red symbols present the number of virions that can pass through 0.1 um filtration after the polyphenol treatment, serial dilution, and vortex. Arrows indicate the detection limit of the one-step RT-qPCR.

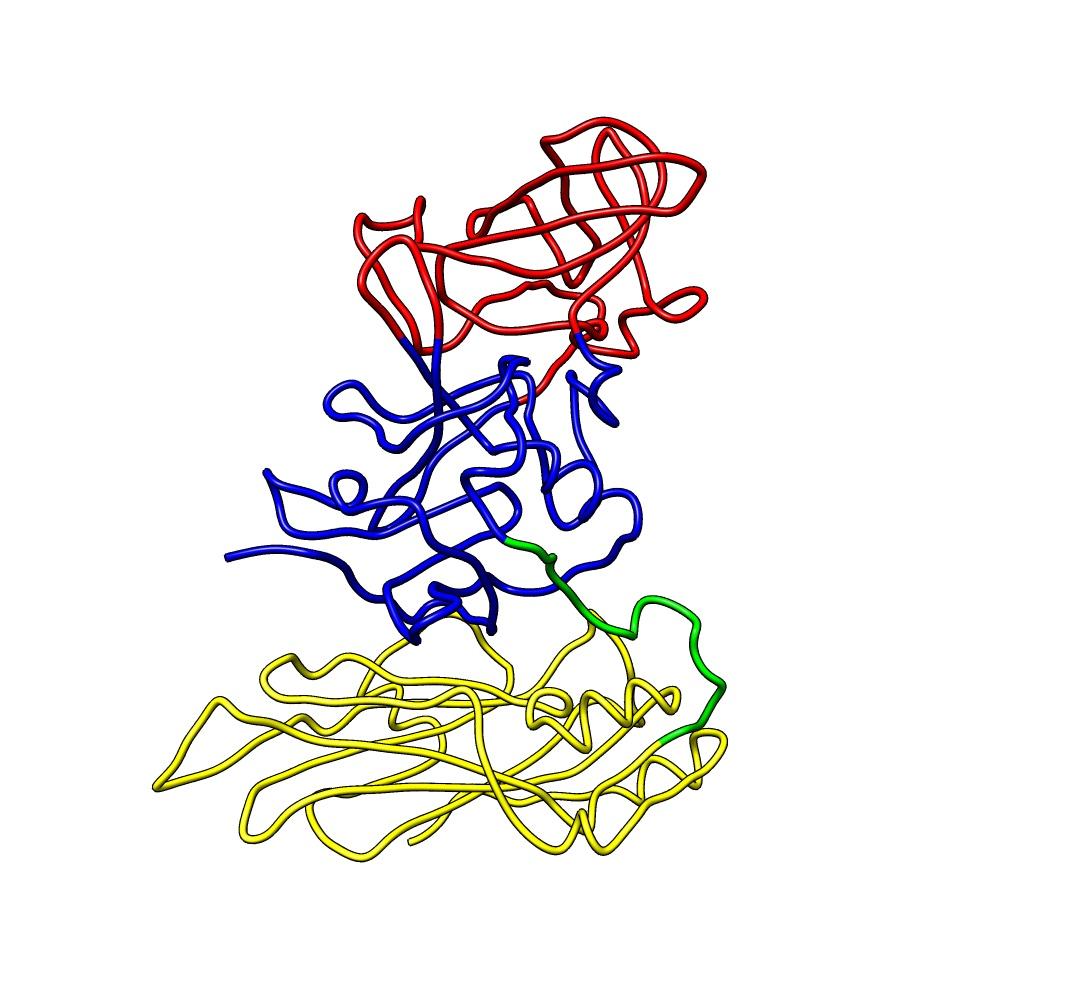

**Supplementary Fig. 2.** VP1 protein structure of human norovirus. Protein information downloaded from the Protein Data Bank (icosahedral asymmetric unit; PDB accession ID: 1IHM) consists of three chains of VP1 proteins, we presented one of the chains. The image was created by Chimera. Each color (Yellow, Green, Blue, and Red) indicated S domain, S-P1 hinge, P1 domain, and P2 domain, respectively.

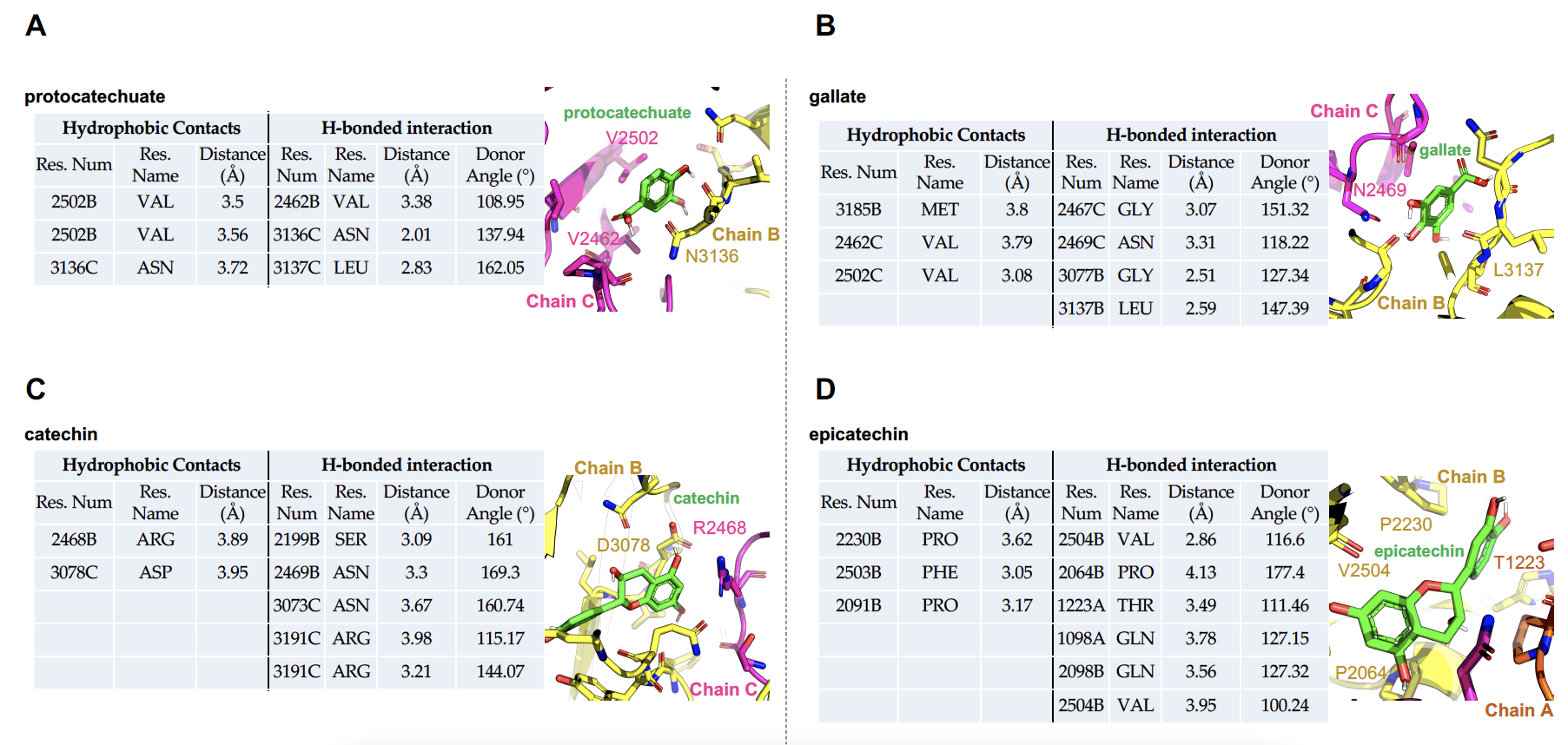

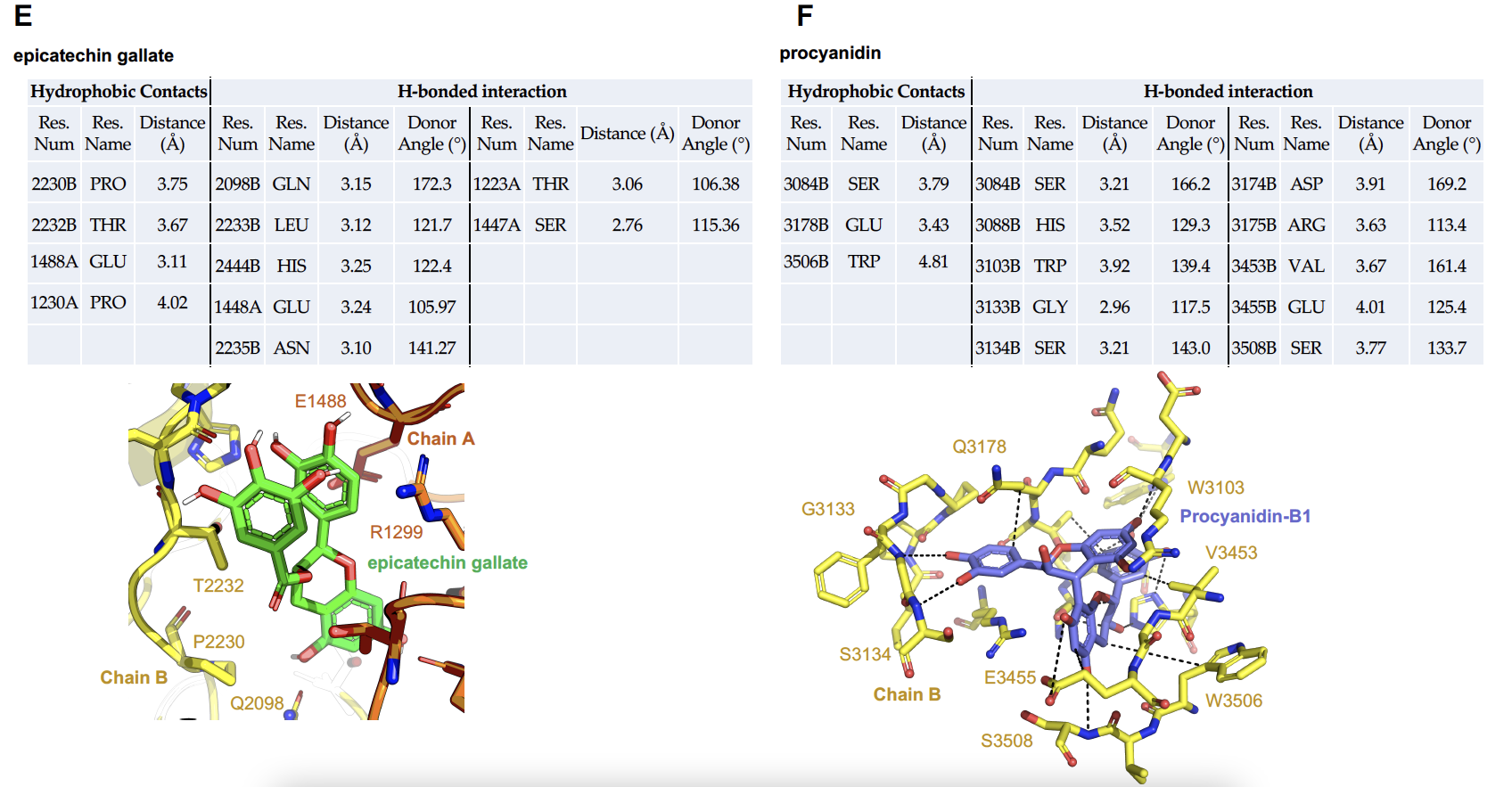

**Supplementary Fig. 3.** The polyphenol-capsid interaction profiles for each polyphenolic compound (A-F) in increasing order of size of polyphenol. For each case, the bound confirmation with the HuNoV capsid is shown and the interacting residues (with details) have been listed alongside. We note that, with an increase in polyphenol size, the number of interactions (both hydrophobic and electrostatic) increases. Additionally, all polyphenols barring the procyanidins bind to interface domains of two chains (between A-copper, B-yellow, or C-magenta). Procyanidins being the largest polyphenol cannot fit into the smaller chain-interface grooves; it is primarily a surface binder as seen from panel (F) - as all of its interactions are limited to only chain B.

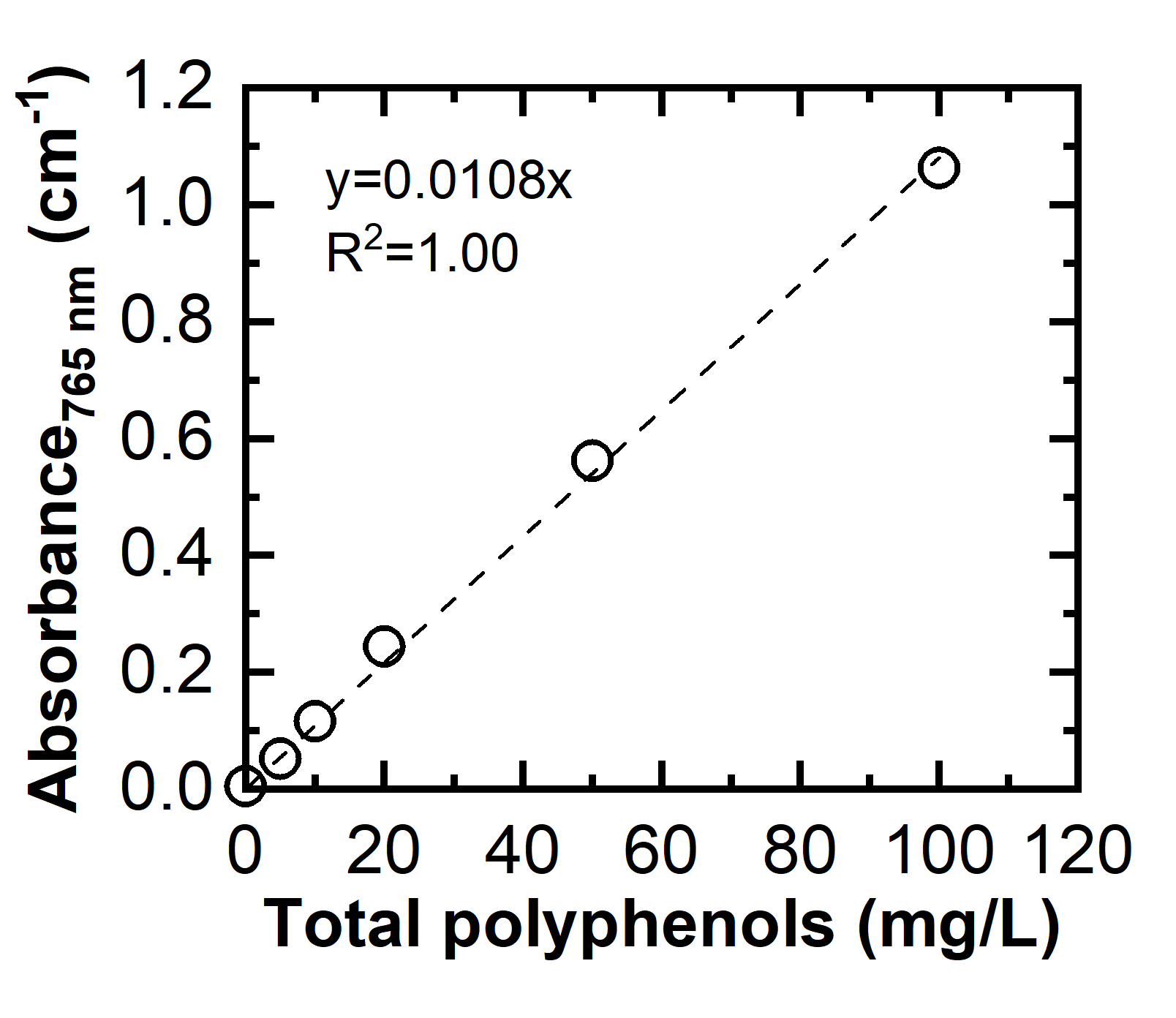

**Supplementary Fig. 4.** A calibration curve of standard solutions for total polyphenols. Gallic acid monohydrate was diluted in distilled water to get the standard TP solutions of 0, 5, 10, 20, 50, and 100 μg/mL. The y-axis shows the absorbance at 765 nm.

**Supplementary Table 2.** Primary polyphenolic compounds observed in the GSE that used for molecular docking simulations

| Category | Polyphenolic compounds | Molecular formula | Theoretical molecular weight (g/mol) | Observed molecular weight (g/mol) |
| --- | --- | --- | --- | --- |
| Monomers flavan-3-ol | Catechin | C_15_H_14_O_6_ | 290.268 | 290.079 |
|  | Epicatechin | C_15_H_14_O_6_ | 290.268 | 290.079 |
|  | Epicatechin gallate | C_22_H_18_O_10_ | 442.372 | 442.090 |
| Dimers flavan-3-ol | Procyanidin B1 | C_30_H_26_O_12_ | 578.520 | 578.143 |
|  | Procyanidin B2 | C_30_H_26_O_12_ | 578.520 | 578.143 |
|  | Procyanidin B3 | C_30_H_26_O_12_ | 578.520 | 578.143 |
|  | Procyanidin B4 | C_30_H_26_O_12_ | 578.520 | 578.143 |
| Phenolic acid | Gallic acid | C_7_H_6_O_5_ | 170.120 | 170.020 |
|  | Protocatechuic acid | C_7_H_6_O_4_ | 154.120 | 154.025 |

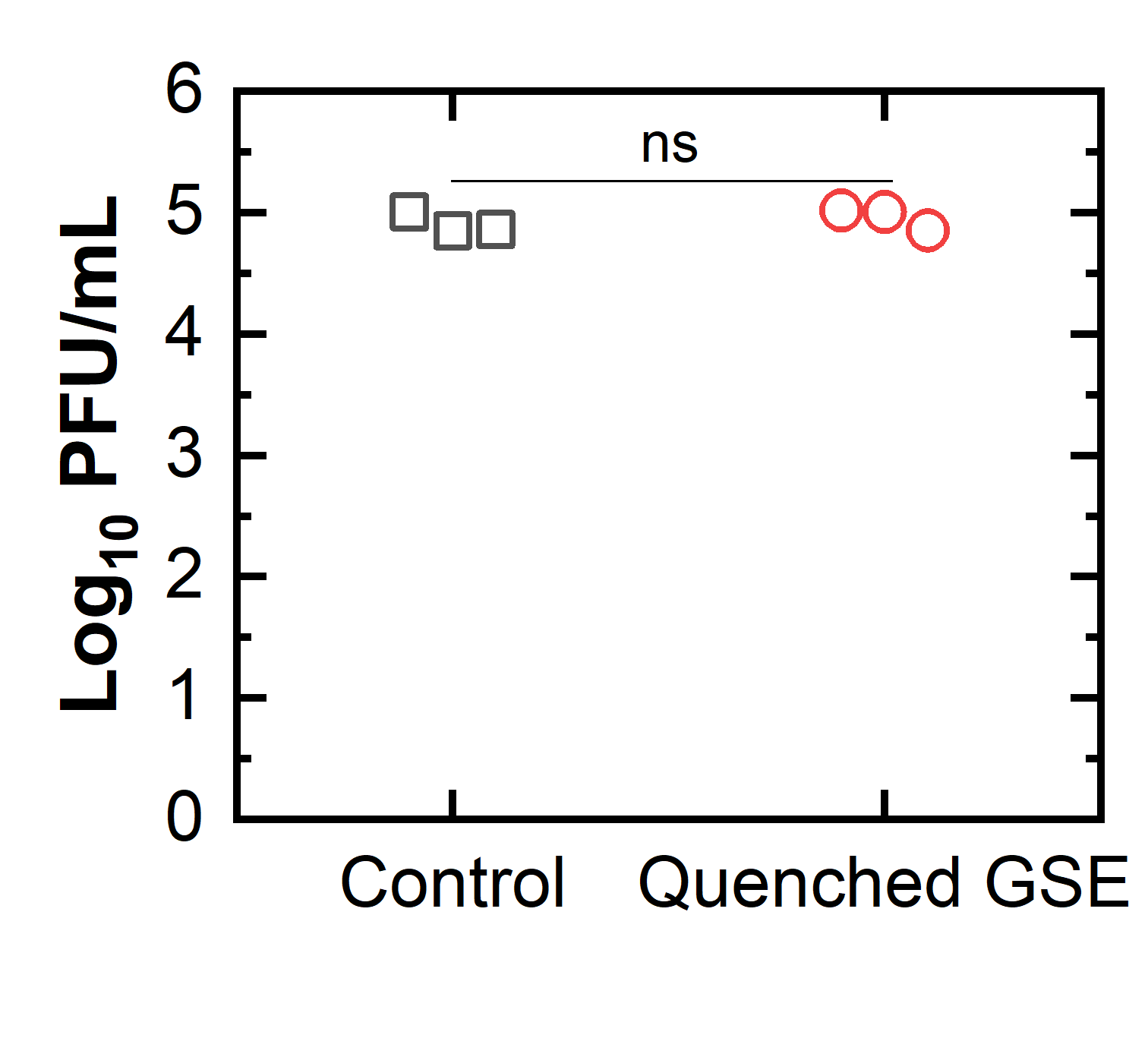

**Supplementary Fig. 5.** Quenching effect of FBS on polyphenolic activity. The “Control” sample was prepared by mixing 10 μL of 1X PBS and 20 μL of FBS followed by adding it to 10 μL of 10^6^ PFU/mL TV. The “Quenched GSE” sample was prepared by quenching 10 μL of 1694 μg/mL total polyphenols by 20 μL of FBS. Then, 10 μL of 10^6^ PFU/mL TV was added to the quenched GSE solution (i.e., the volume ratio of TV, GSE, and FBS was 1:1:2). The y-axis indicates TV infectivity determined by the plaque assay. Mann-Whitney test determined the infectivities of “Control” and “Quenched GSE” samples were not significantly different (p>0.05). This result means that FBS can quench the antiviral activity of 847 μg/mL total polyphenols in a volume ratio of 1:1, the highest total polyphenol concentration in this research.

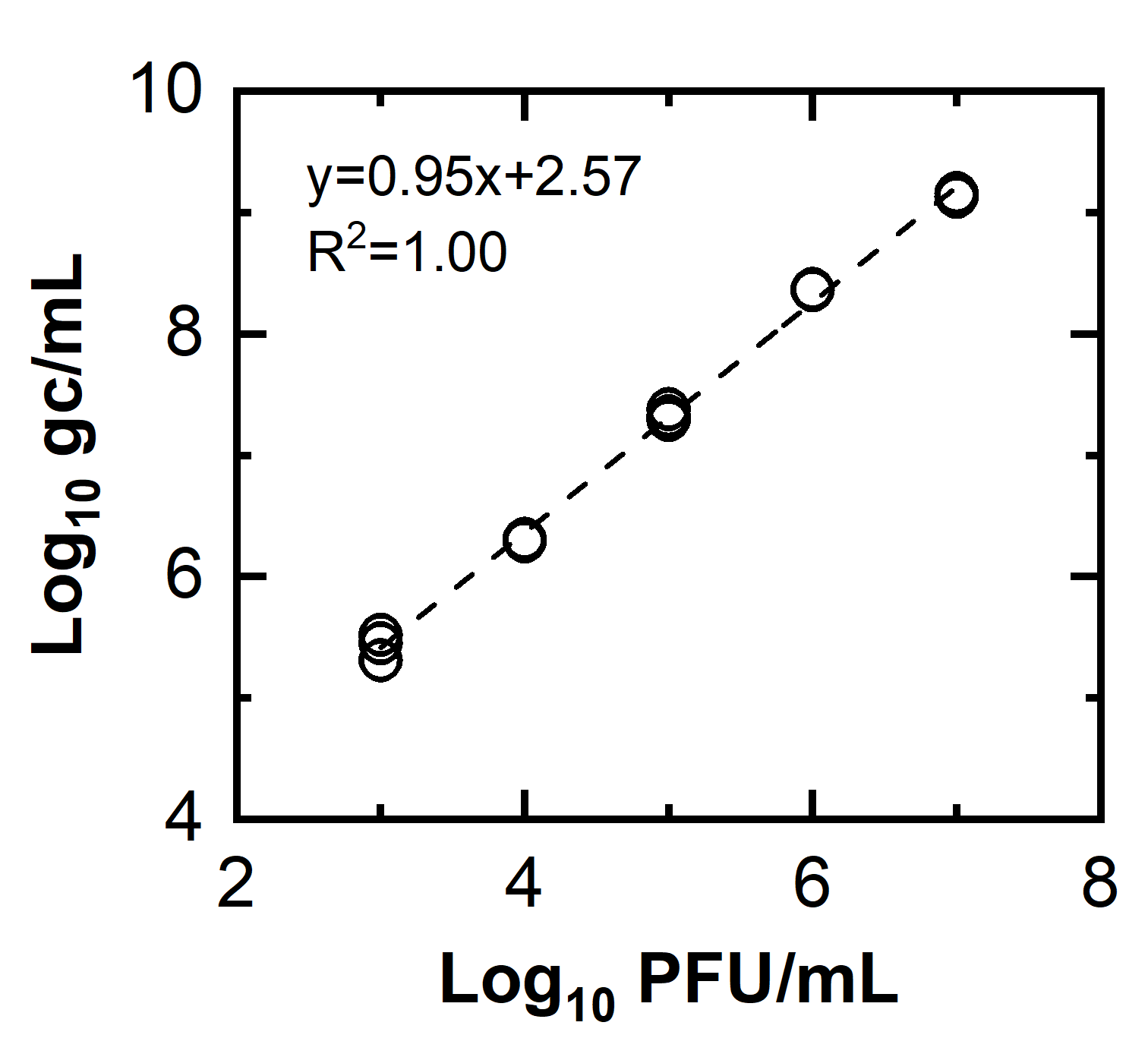

**Supplementary Fig. 6.** A calibration curve for the aggregation assay. The x-axis indicates infectivity of the initial TV and the y-axis shows the gene copy number determined by the one-step RT-qPCR assay. The reliable range of the aggregation assay was determined between 10^3^ and 10^7^ PFU/mL.

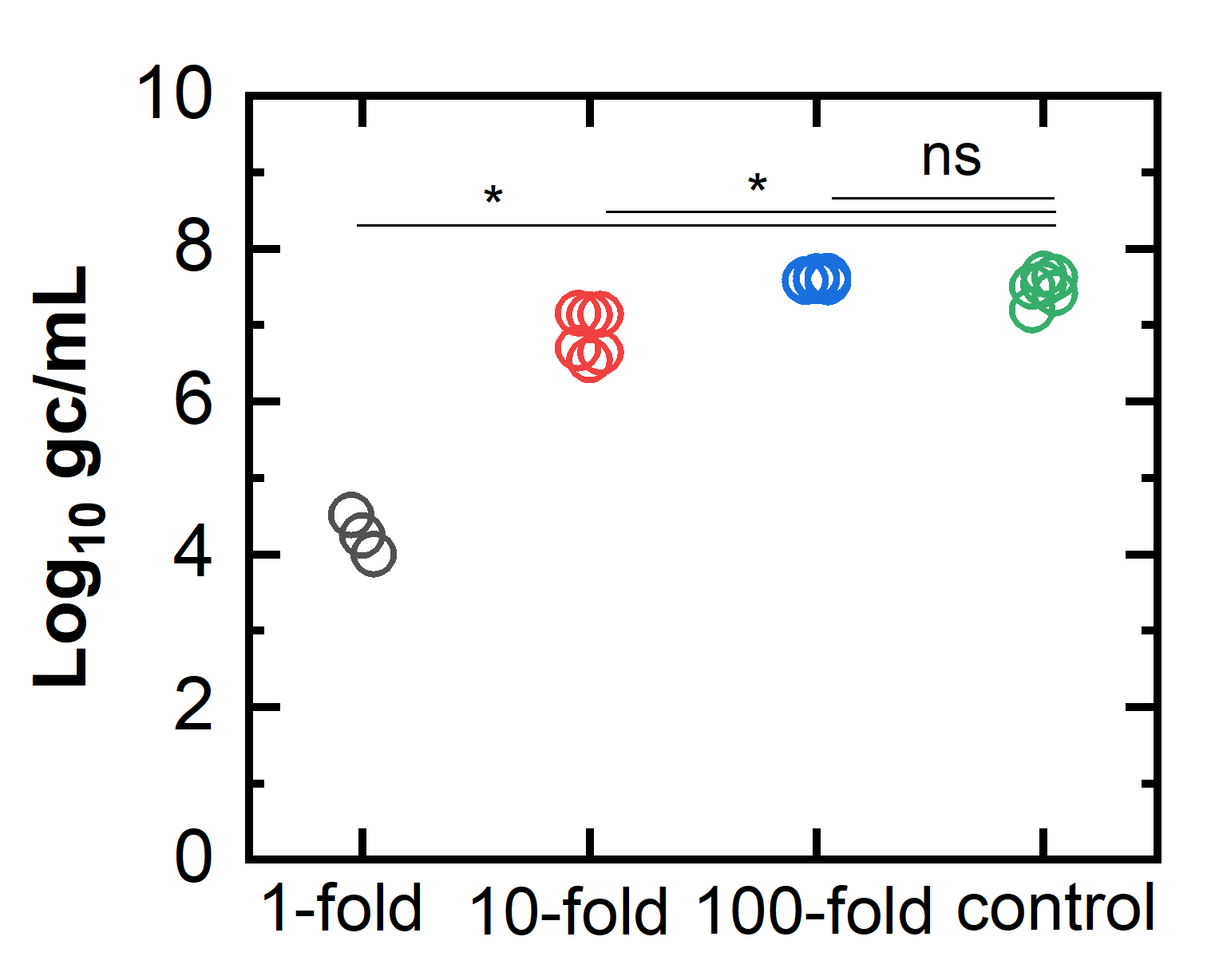

**Supplementary Fig. 7.** Inhibition effect of polyphenols on the extracted genomic RNA. The y-axis represents the RT-qPCR assay results and the x-axis indicates sample preparation methods: the 1X means that 25 μL of 10^7^ PFU/mL TV was mixed with 25 μL of 1694 μg/mL GSE and 50 μL FBS was added to the mixture. The 10-fold and 100-fold indicate 10- and 100-fold dilution of the 1X solution, respectively. Control meant 25 μL PBS replaced the 25 μL GSE in the 1X solution (i.e., TV:PBS:FBS=25:25:50 uL). Mann-Whitney test analyzed the statistical differences between the two groups. Both 1X and 10X showed significant decreases in copy number/mL while 100-fold dilution did not present a significant difference compared to the control, indicating the 100-fold dilution was required for the RT qPCR assay to prevent the extracted RNA from being inhibited by GSE.

**Supplementary Table 3.** Information for RT-qPCR conditions and primers

| **process** | **Primer name** | **Sequence^1)^ (5’-3’)** | **Position in the genome** | **Amplicon length (bp)** | **Reaction conditions** |
| --- | --- | --- | --- | --- | --- |
| RT-qPCR | TV-NSP1-qPCR-F | GTGCGCATCCTTGAGACAAT | 879-899 | 132^2)^ | 50°C for 10 min and 95°C for 1 min, followed by 40 cycles of (95°C for 10 s, 60°C for 30 sec) |
|  | TV-NSP1-qPCR-R | TTGGAGCCGGGTAGAAACAT | 991-1011 |  |  |

1) Primer pair specificity was checked by the primer-blast tool (National Center of Biotechnology Information). Each pair of primers were blasted with Rhesus monkey (taxid:9544) which was the host organism. As a result, we confirmed that our primers did not target any sequences of the MA104 cells.
2) The sequence of standard sample for the NSP gene of Tulane virus (Integrated DNA technologies, USA):

5’-AGAATTGGACCGAATTTGGCACACACTCAGAATTTGGTGTGCGCATCCTTGAGACAATAACAGGCACAATACCCCCTTGGAAACCTCACCAGGAATCAATATCTGAAGTTCTGGACGACCTCACACACGGTAAAGTCCAAACAGGTGATGATGTTTCTACCCGGCTCCAAAGGTTGAGCGACACTATCAAAGATCTGAGTGTCATGGCTTGTGATCCCTCTGCACCGCCCGAAGTTGCGC-3’ - (GenBank accession number: EU391643).

**Supplementary Table 4.** Checklist for MIQE and location of relevant information in this study

| **Item to check** | **Location** |
| --- | --- |
| **1. Experimental design** | |
| Definition of experimental and control groups | Materials and Methods |
| Number within each group | Materials and Methods; Each figure |
| **2. Sample** | |
| Description | Materials and Methods |
| Volume/mass of sample processed | Materials and Methods |
| Processing procedure | Materials and Methods |
| Sample storage conditions and duration | Materials and Methods |
| **3. Nucleic acid extraction** | |
| Procedure and/or instrumentation | Materials and Methods |
| Name of kit and details of any modifications | Materials and Methods |
| Contamination assessment (DNA or RNA) | Materials and Methods |
| Nucleic acid quantification | Materials and Methods |
| Instrument and method | Materials and Methods |
| Inhibition testing (C_q_ dilutions, spike, or other) | Supplementary Fig. 7 |
| **4. Reverse transcription** | |
| Complete reaction conditions | Materials and Methods; Supplementary Table 3 |
| Amount of RNA and reaction volume | Materials and Methods |
| Reverse transcriptase and concentration | Materials and Methods |
| Temperature and time | Supplementary Table 3 |
| Manufacturer of reagents and catalog numbers | Materials and Methods |
| **5. qPCR target information** | |
| Gene symbol | Supplementary Table 3 |
| Sequence accession number | Supplementary Table 3 |
| Location of amplicon | Supplementary Table 3 |
| Amplicon length | Supplementary Table 3 |
| In silico specificity screen (BLAST, and so on) | Supplementary Table 3 |
| **6. qPCR oligonucleotides** | |
| Primer sequences | Supplementary Table 3 |
| Manufacturer of oligonucleotides | Supplementary Table 3 |
| **7. qPCR protocol** | |
| Complete reaction conditions | Materials and Methods; Supplementary Table 3 |
| Reaction volume and amount of DNA | Materials and Methods |
| Primer (probe) concentrations | Materials and Methods |
| Polymerase identity and concentration | Materials and Methods |
| Buffer/kit identity and manufacturer | Materials and Methods |
| Manufacturer of plates/tubes and catalog number | Materials and Methods |
| Complete thermocycling parameters | Materials and Methods; Supplementary Table 3 |
| Manufacturer of qPCR instrument | Materials and Methods |
| **8. qPCR validation** | |
| Specificity (gel, sequence, melt, or digest) | Materials and Methods |
| For SYBR Green I, C_q_ of the NTC | Materials and Methods |
| Calibration curves with slope and *y-intercept* | Materials and Methods; Supplementary Fig. 6 |
| PCR efficiency calculated from slope | Materials and Methods |
| *r*2 of calibration curves | Materials and Methods |
| Linear dynamic range | Materials and Methods |
| C_q_ variation at LOD | Materials and Methods |
| Evidence for LOD | Materials and Methods |
| **9. Data analysis** | |
| qPCR analysis program (source, version) | Materials and Methods |
| Method of C_q_ determination | Materials and Methods |
| Outlier identification and disposition | Materials and Methods |
| Results for NTCs | Materials and Methods |
| Description of normalization method | Materials and Methods |
| Number and concordance of biological replicates | Materials and Methods; each Figure |
| Number and stage of technical replicates | Materials and Methods |
| Repeatability (intraassay variation) | Materials and Methods |
| Statistical methods for results significance | Each figure |
| Software (source, version) | Materials and Methods |
| Data transparency | Raw data available upon request |

**References**

1.	Gerba CP, Betancourt WQ. 2017. Viral Aggregation: Impact on Virus Behavior in the Environment. Environ Sci Technol 51:7318–7325.

2.	Mattle MJ, Kohn T. 2012. Inactivation and tailing during UV254 disinfection of viruses: Contributions of viral aggregation, light shielding within viral aggregates, and recombination. Environ Sci Technol 46:10022–10030.

3.	Floyd R, Sharp DG. 1977. Aggregation of poliovirus and reovirus by dilution in water. Appl Environ Microbiol 33:159–167.

4.	Kahler AM, Cromeans TL, Metcalfe MG, Humphrey CD, Hill VR. 2016. Aggregation of Adenovirus 2 in Source Water and Impacts on Disinfection by Chlorine. Food Environ Virol 8:148–155.

5.	Yu G, Zhang D, Guo F, Tan M, Jiang X, Jiang W. 2013. Cryo-EM Structure of a Novel Calicivirus, Tulane Virus. PLoS One 8.
